# Interactions between TULP3 tubby domain cargo site and ARL13B amphipathic helix promote lipidated protein transport to cilia

**DOI:** 10.1101/2021.05.25.445488

**Authors:** Vivek Reddy Palicharla, Sun-Hee Hwang, Bandarigoda N. Somatilaka, Hemant B. Badgandi, Emilie Legué, Vanna M. Tran, Jeffrey B. Woodruff, Karel F. Liem, Saikat Mukhopadhyay

## Abstract

The tubby family protein–TULP3 coordinates with the intraflagellar transport complex-A (IFT-A) in trafficking certain transmembrane proteins to cilia. These transmembrane cargoes have short motifs that are necessary and sufficient for TULP3-mediated trafficking. However, whether TULP3 regulates trafficking of membrane-associated proteins is not well understood. Here we show that TULP3 is required for transport of the atypical GTPase ARL13B into cilia, and for ciliary enrichment of ARL13B-dependent farnesylated and myristoylated proteins. ARL13B transport requires TULP3 binding to IFT-A core but not to phosphoinositides, unlike transmembrane cargo transport that requires binding to both by TULP3. A conserved lysine in TULP3’s tubby domain mediates direct ARL13B binding and trafficking of lipidated and transmembrane cargoes. An N-terminal amphipathic helix in ARL13B flanking the palmitoylation site mediates binding to TULP3 and directs trafficking to cilia even in absence of palmitoylation and RVxP sorting motif. Therefore, TULP3 transports transmembrane proteins and ARL13B into cilia by capture of short sequences through a shared tubby domain site.

## Introduction

The primary cilium is a microtubule based dynamic cellular appendage that is templated from the mother centriole of the centrosome–the basal body (Anvarian et al., 2019). Cilia are present in multiple cell types and organs and can transduce cellular response to extracellular signals. These signals include hedgehog morphogens that regulate multiple developmental and regenerative programs (Kopinke et al., 2020). Signaling outputs from cilia also maintain renal tubular homeostasis preventing cystogenesis (Ma, 2021). The ciliary membrane has a distinct protein and lipid composition despite being contiguous with the plasma membrane (Nachury and Mick, 2019). Multiple transmembrane (Hilgendorf et al., 2016) and lipidated proteins are preferentially associated with the ciliary membrane. The lipidated proteins include palmitoylated proteins such as ARL13B (Caspary et al., 2007; Duldulao et al., 2009), farnesylated proteins such as INPP5E (Bielas et al., 2009) and myristoylated proteins such as NPHP3 (Shiba et al., 2010). Deciphering mechanisms underlying ciliary membrane compartmentalization is necessary to understand how signaling at and by cilia contributes to morpho-phenotypic outcomes.

ARL13B is an atypical GTPase that is highly enriched in the primary cilium. Mutations in *ARL13B* in humans are associated with classic Joubert syndrome, a ciliopathy associated with cerebellar vermis hypoplasia, hypotonia, intellectual disability (Cantagrel et al., 2008) and occasionally accompanied with renal cysts, retinal impairment and obesity (Thomas et al., 2015). ARL13B acts as a GEF for the GTPase ARL3, converting inactive ARL3-GDP to active ARL3-GTP (Gotthardt et al., 2015; Ivanova et al., 2017). ARL3-GTP promotes ciliary release of multiple lipidated cargoes including farnesylated and myristoylated proteins from their respective carrier proteins–Pde6*δ* (Humbert et al., 2012) and Unc119b (Wright et al., 2011). Given that ARL13B potentially regulates levels of multiple lipidated proteins inside cilia, it seems logical that it affects ciliary signaling. However, the role of trafficking of ARL13B into cilia is tissue specific. The zebrafish *arl13b* (*scorpion*) allele has nephric duct dilatation phenotypes, and analysis of phenotypic rescue using *arl13b* variants in this model suggest that ciliary localization is essential for *in vivo* function of Arl13b (Duldulao et al., 2009; Sun et al., 2004). However, a mouse knock-in at the endogenous locus, which prevents the mutant protein from localizing to cilia (*Arl13b^V358A^*), is not affected in neural tube patterning (Gigante et al., 2020), unlike the null *Arl13b^hnn^* mutant where neural tube patterning is affected (Caspary et al., 2007). A precise understanding of mechanisms trafficking ARL13B to cilia could perhaps shed light on the differences observed in the role of ciliary trafficking of ARL13B in different contexts.

A definitive ciliary targeting mechanism for ARL13B has been lacking, mostly because of contrasting results with regards to ciliary targeting sequences that are necessary but not sufficient for trafficking (see supplemental text). One motif that has stood out as being necessary for ciliary localization is the RVxP motif in the C-terminus (Mariani et al., 2016). Other studies have reported cellular aggregates upon deletion of the RVxP motif (Cevik et al., 2013; Nozaki et al., 2017), along with proximal to distal ciliary mislocalization in *C. elegans* phasmid cilia (Cevik et al., 2013). The RVxP motif was initially discovered as a C-terminal motif involved in post-Golgi trafficking of Rhodopsin (Deretic et al., 2005; Wang et al., 2012) and other proteins in the secretory pathway, including the olfactory CNGB1B subunit (Jenkins et al., 2006), and Polycystin-1 and Polycystin-2 channels (Geng et al., 2006; Ward et al., 2011). ARL13B is a cytosolic protein that is palmitoylated, and the RVxP motif is not strictly at the end of the protein (unlike within 10 aa of the C-terminus in other secretory pathway proteins). The interpretations of experiments in different settings are further complicated by ciliary localization assays of ARL13B mutants performed in the wildtype (Cevik et al., 2010; Cevik et al., 2013; Hori et al., 2008; Nozaki et al., 2017) or null background (Duldulao et al., 2009; Larkins et al., 2011; Mariani et al., 2016). Experiments in *C. elegans* have also demonstrated dominant negative roles of mutants (Cevik et al., 2010; Cevik et al., 2013), suggesting mutual interactions in compromising wild-type protein function. Self-association of ARL13B through its N-terminal domain has been reported (Hori et al., 2008) that could explain some of the above dominant negative effects, although others have reported lack of dimerization of the purified GTPase domain (Miertzschke et al., 2014). Some of the ARL13B mutants, such as deletion of the RVxP motif or lack of the coiled coil, form cellular aggregates and could be indirectly affecting ciliary targeting (Cevik et al., 2013; Nozaki et al., 2017). In sum, the mechanism of ARL13B trafficking by the RVxP motif and in different contexts has remained unclear.

The tubby family protein, TULP3, functions in coordination with the intraflagellar transport complex-A (IFT-A) to determine ciliary trafficking of transmembrane proteins without affecting their extraciliary pools or disrupting cilia (Mukhopadhyay et al., 2010). These cargoes include GPCRs and polycystin channels (Badgandi et al., 2017). We proposed three steps during TULP3 mediated trafficking to cilia that consist of 1) capturing of the membrane localized cargoes by TULP3’s C-terminal tubby domain in a PI(4,5)P_2_-dependent manner, 2) trafficking into the primary cilium via IFT-A core binding to the N-terminus of TULP3, and 3) release of GPCR cargoes from TULP3 into the PI(4,5)P_2_ deficient ciliary membrane. TULP3 and the related but brain-specific Tubby are now established as the major conduit for trafficking of multiple types of membrane cargoes to mammalian cilia (Barbeito et al., 2020; Chavez et al., 2015; Dateyama et al., 2019; Garcia-Gonzalo et al., 2015; Hilgendorf et al., 2019; Hirano et al., 2017; Loktev and Jackson, 2013; Mukhopadhyay et al., 2017; Sun et al., 2012; Wu et al., 2020).

In line with a role of TULP3 in trafficking multiple cargoes, TULP3 and IFT-A have been implicated in neural tube patterning (Legue and Liem, 2020; Liem et al., 2012; Mukhopadhyay et al., 2013; Norman et al., 2009; Qin et al., 2011), renal cystogenesis (Hwang et al., 2019; Legue and Liem, 2019; Wang et al., 2020) and adipogenesis (Hilgendorf et al., 2019; Wu et al., 2020). In some cases, such as neural tube development (Mukhopadhyay et al., 2013) or adipogenesis (Hilgendorf et al., 2019), the relevant TULP3-regulated cargoes are likely to be GPCRs, but for other tissues such as renal cystogenesis, the TULP3 cargoes have not been established. We recently showed that kidney collecting duct specific embryonic or juvenile-onset *Tulp3* knockout (ko) in mice results in postnatal kidney cystogenesis, whereas adult-onset loss causes delayed cystogenesis (Hwang et al., 2019; Legue and Liem, 2019). Interestingly, we observed that ARL13B pools in cilia of *Tulp3* conditional ko (cko) collecting ducts were depleted prior to cyst formation in juveniles and in adults, indicating a role for TULP3 in ARL13B ciliary localization (Hwang et al., 2019; Legue and Liem, 2019). However, the direct role of TULP3 in trafficking ARL13B to cilia and relevance of any such trafficking mechanisms have remained unknown.

Our earlier studies showed that for almost all TULP3 dependent transmembrane cargoes, the ciliary targeting motifs were short peptide sequences. When fused to membrane anchors, these sequences were in proximity to TULP3. These motifs were both necessary and sufficient for ciliary trafficking (Badgandi et al., 2017). Here, we show that TULP3 is required for ciliary trafficking of ARL13B, and enrichment of ARL13B-dependent farnesylated and myristoylated proteins in cilia. We identify an amphipathic helix in the N-terminus of ARL13B to mediate binding to TULP3 and in determining ciliary localization of ARL13B. We identify the corresponding site in the tubby domain of TULP3 that mediates ciliary trafficking of both ARL13B and transmembrane cargoes. Thus, TULP3, by binding to short sequences through shared tubby domain interfaces determines levels of transmembrane cargoes and ARL13B-dependent lipidated cargoes in cilia.

## Results

### TULP3 is required for ciliary trafficking of ARL13B

We recently observed that cilia in kidney collecting ducts deleted for *Tulp3* lacked Arl13b (Hwang et al., 2019; Legue and Liem, 2019). To further investigate the role of Tulp3 in ciliary trafficking of Arl13b in different contexts, we first tested Arl13b ciliary trafficking in mouse embryonic fibroblasts (MEFs) obtained from wild type (WT) and *Tulp3* knockout (ko) mice (Legue and Liem, 2019). Arl13b was primarily localized to cilia in wild type (WT) MEFs whereas ciliary localization was abrogated in *Tulp3* ko MEFs, suggesting that Arl13b ciliary localization is Tulp3 dependent (**Figure 1A, 1B**). Ciliary localization of Gpr161, a known GPCR cargo for Tulp3 (Mukhopadhyay et al., 2013), was also lost in *Tulp3* ko MEFs as expected (**Figure 1B, S1A**). *Tulp3* ko MEFs had no observable defects in ciliation with respect to number (**Figure 1B**) but showed a small reduction in length (**Figure 1C**). We further generated *Tulp3* ko 3T3-L1 preadipocyte and NIH 3T3 cells using CRISPR/Cas9 technology (**Figure S1B**). We observed that Arl13b ciliary localization was inhibited in *Tulp3* ko 3T3-L1 and NIH 3T3 cells (**Figure 1D-1E, Figure S1C-D**), confirming that Tulp3 is required for Arl13b ciliary localization. While ciliation with respect to number was unaffected (**Figure 1E**), a small reduction in ciliary length was observed in *Tulp3* ko 3T3-L1 cells (**Figure 1F**). Smoothened, a Tulp3 independent cargo (Mukhopadhyay et al., 2010), was trafficked to *Tulp3* ko 3T3-L1 cells upon treatment with a Smoothened agonist SAG (**Figure S1E-F**), ruling out generalized effects on localization from defects in transition zone (Chih et al., 2011).

**Figure 1.**
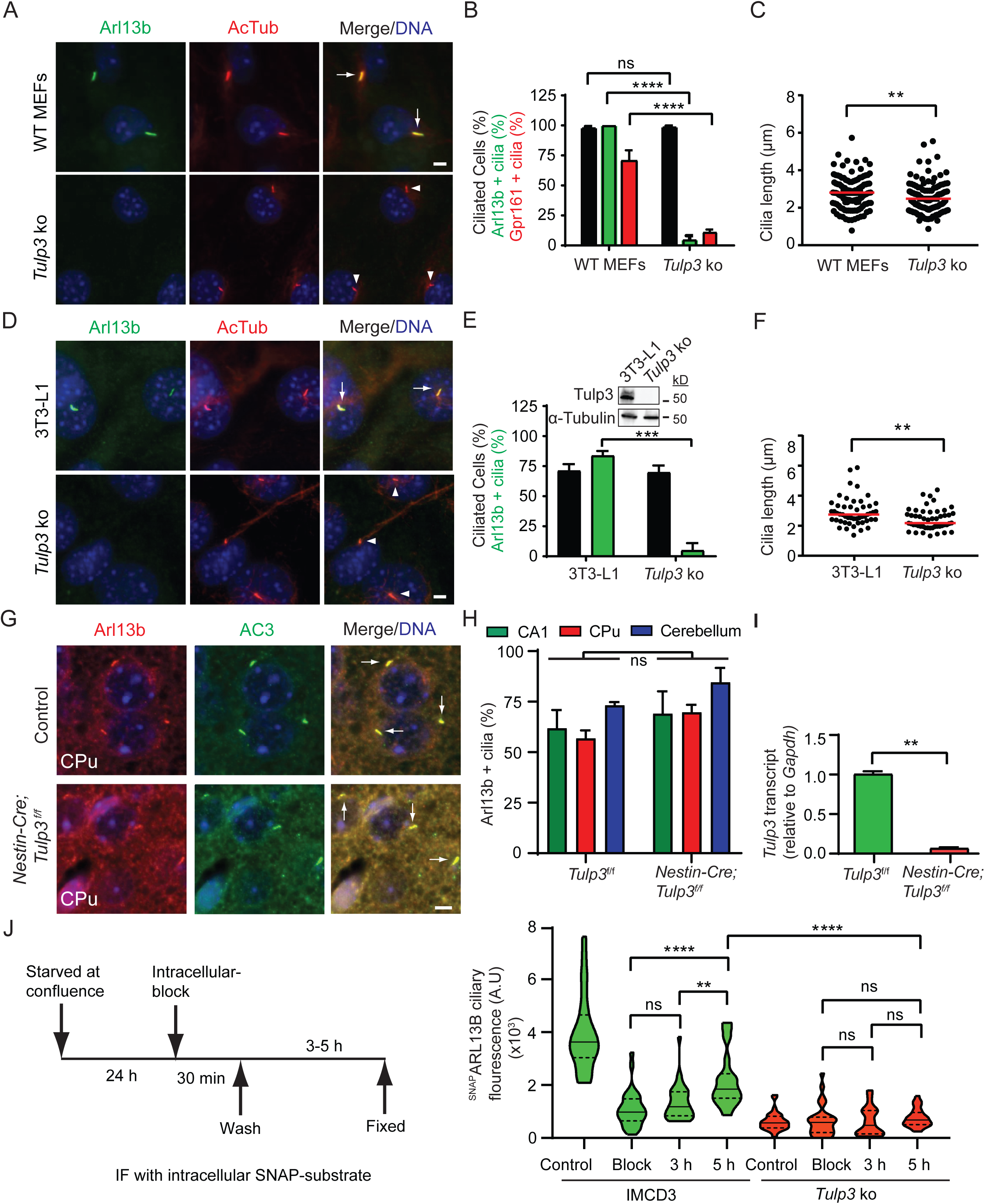
Tulp3 determines ciliary trafficking of Arl13b. **(A)** MEFs from wild type or *Tulp3* knockout (ko) mice were serum starved upon confluence for 24 h before fixation. Fixed cells were immunostained for Arl13b (green) or Gpr161 (shown in Figure S1) along with acetylated tubulin (AcTub, red) and counterstained for DNA (blue). **(B)** Arl13b (from A) or Gpr161 (from Figure S1) positive cilia were counted from two experiments, and total counted cells are >200 for each condition. Data represent mean ± SD. **(C)** Cilia length was measured from wild type or *Tulp3* ko MEFs by immunostaining for acetylated tubulin. Number of cilia measured for length is >100 for each condition. **(D)** Wildtype and *Tulp3* ko 3T3-L1 preadipocyte cells grown to confluency and cultured further for 72 h to promote ciliation before fixing. The fixed cells were immunostained for Arl13b (green) along with acetylated tubulin (red) and counterstained for DNA. **(E)** Arl13b positive cilia from D were counted from three experiments, and total counted cells are >200 for each condition. Data represent mean ± SD. Inset shows immunoblot of wild type or *Tulp3* ko 3T3L1 cell lysates with Tulp3 and Tubulin antibodies. **(F)** Cilia length was measured from wild type or *Tulp3* ko 3T3L1 cells by by immunostaining for acetylated tubulin. n>50 for each condition. **(G)** Brain sections from caudate-putamen (CPu) and CA1 regions in hippocampus from either control or *Nestin-Cre*; *Tulp3*^f/f^ mice at P7 were immunostained for Arl13b and AC3 and counterstained for DNA. **(H)** Arl13b positive cilia similar to (G) were counted from control or conditional knockout (cko) at P7. Total counted cilia are >400 for caudate-putamen regions, 50-250 for CA1 regions, and >650 for cerebellums for each mouse from one control and two cko mice. Data represent mean ± SD. **(I)** *Tulp3* transcript levels relative to *Gapdh* were measured from control or Nestin Cre; *Tulp3*^f/f^ mice. Data represents mean ± SEM from two data points from one mouse. **(J)** Wild type and *Tulp3* ko IMCD3 cells stably expressing ^GFP-SNAP^ARL13B were starved for a total of 36 h after reaching confluence. After blocking intracellular pools of ^GFP-SNAP^ARL13B using BG-Block, cells were washed and later fixed at indicated times. “Block” refers to cells immediately fixed after blocking. Control refers to cells untreated with BG-block. Newly trafficked SNAP-tagged proteins in cilia in BG-block treated cells were tracked by immunofluorescence using fluorescent SNAP substrate (TMR-Star) and acetylated tubulin along with counterstaining for DNA. Images in **Figure S1C**. Violin plots of ciliary intensities of TMR-Star from n>30 cells for each condition. Scale, 5 μm; ****, p<0.0001; ***, p<0.001; **, p<0.01; ns, not significant. Arrows indicate cilia positive for the indicated proteins while arrowheads indicate negative cilia. See also **Figure S1** and **Figure S2**.

Tulp3 belongs to tubby protein family consisting of four other members, Tubby, Tulp1, Tulp2 and Tulp4 (Mukhopadhyay and Jackson, 2011). Tulp3 is ubiquitously expressed, while Tubby is expressed only in brain and retina (Mukhopadhyay and Jackson, 2011). Tubby also binds to IFT-A, but to a lesser extent (Badgandi et al., 2017; Mukhopadhyay et al., 2010) and regulates ciliary targeting of GPCRs (Badgandi et al., 2017; Loktev and Jackson, 2013; Sun et al., 2012). Unlike kidney tubules (Hwang et al., 2019; Legue and Liem, 2019), we observed that ciliary localization of Arl13b was unaffected in *Nestin-Cre*; *Tulp3^f/f^* brain regions at P7 (caudate putamen/striatum, dentate gyrus and CA1 region of the hippocampus) (**Figure 1G-H, Figure S2A-B**) and in postnatal cerebellum lobes at P7 (**Figure 1H**). RNA transcripts of *Tulp3* were reduced in *Tulp3* cko brain (**Figure 1I**). Arl13b has also been reported to be unaffected in cilia of *Tulp3* ko mouse neural tube (Ferent et al., 2019). Therefore, Arl13b is unaffected in *Tulp3* deletions in brain and neural tube, likely from redundancy with Tubby.

### TULP3 determines entry of ARL13B into cilia

Steady state levels of a protein in cilia can be regulated at multiple steps, including trafficking to cilia, removal from the compartment, or additional steps in cargo recycling (Mukhopadhyay et al., 2017). We previously demonstrated that Tulp3 determined entry of Gpr161 into cilia, rather than inhibiting removal from the compartment (Badgandi et al., 2017). Therefore, we tested the direct role of Tulp3 in Arl13b entry into cilia. We tested the dynamics of ciliary entry of stably expressed ARL13B that was tagged at the N-terminus with GFP-SNAP (^GFP-SNAP^ARL13B) in control and *Tulp3* ko inner medullary collecting duct (IMCD3) cells (**Figure 1J**), generated using CRISPR/Cas9 technology (**Figure S1B**). After blocking intracellular pools of GFP-SNAP tagged ARL13B using non fluorescent SNAP substrate, we tracked accumulation of newly trafficked SNAP-tagged proteins in cilia by immunofluorescence using fluorescent SNAP substrate (TMR-Star) at different time points. We detected an increase of up to ∼50% of mean steady state SNAP-labeled pools by 5 h in cilia of control cells. Such increase was strongly reduced in *Tulp3* ko cells, suggesting inhibition of ciliary entry of ARL13B (**Figure 1J, Figure S2C**).

### TULP3 is required for enrichment of both farnesylated and myristoylated proteins in cilia

Arl13b acts as a GEF for Arl3 converting inactive Arl3-GDP to active Arl3-GTP (Gotthardt et al., 2015; Ivanova et al., 2017). Arl3 once activated promotes ciliary release of multiple lipidated cargoes including farnesylated proteins such as Inpp5e and myristoylated proteins such as Nphp3 and Cystin1 (Cys1) from their respective carrier proteins Pde6*δ* (Humbert et al., 2012) and Unc119b (Wright et al., 2011), respectively (**Figure 2A**). As we observed that Arl13b ciliary trafficking was dependent on Tulp3, we asked if Tulp3 is also required for ciliary trafficking of Arl13b/Arl3 dependent cargoes. To answer this question, we stably expressed GFP-Stag (LAP) tagged INPP5E, NPHP3 (1-203 aa) or CYS1 (**Figure 2B, 2D**) in control or *Tulp3* ko IMCD3 cells (**Figure S1B**). Compared to control, *Tulp3* ko IMCD3 cells showed loss of endogenous Arl13b and ^GFP^INPP5E ciliary localization (**Figure 2B-C**). Importantly, ciliary trafficking of Arl13b and ^GFP^INPP5E were restored upon stably expressing ^Myc^TULP3 (**Figure 2B-C**) ruling out effects from nonspecific deletions in the genome. Total cellular levels of endogenous Arl13b protein were unaffected by immunoblotting in *Tulp3* ko (**Figure 2C**). Localization of NPHP3^GFP^ and CYS1^GFP^ were reduced in intensities but were partially retained in cilia when quantified as cilia positive for respective proteins irrespective of decreased intensities (**Figure 2D-F**). Lkb1 is a farnesylated protein that is targeted to cilia by the pseudokinase STRAD*β* (Mick et al., 2015). Together, they form a heterotrimer complex with the scaffolding protein MO25 (Zeqiraj et al., 2009). Lkb1 trafficking is therefore likely to be independent of Arl3. In agreement with this, ciliary localization of endogenous Lkb1 was only modestly reduced in *Tulp3* ko with no significant decrease in ciliary intensities (**Figure 2G-I**), suggesting Lkb1 trafficking to be independent of Tulp3. We also observed a small decrease in ciliary length upon *Tulp3* ko in IMCD3 cells that was partially rescued by stably expressing ^Myc^TULP3 (**Figure 2J**). Thus, Tulp3 determines ciliary localization of Arl13b and Arl13b dependent lipidated cargoes.

**Figure 2.**
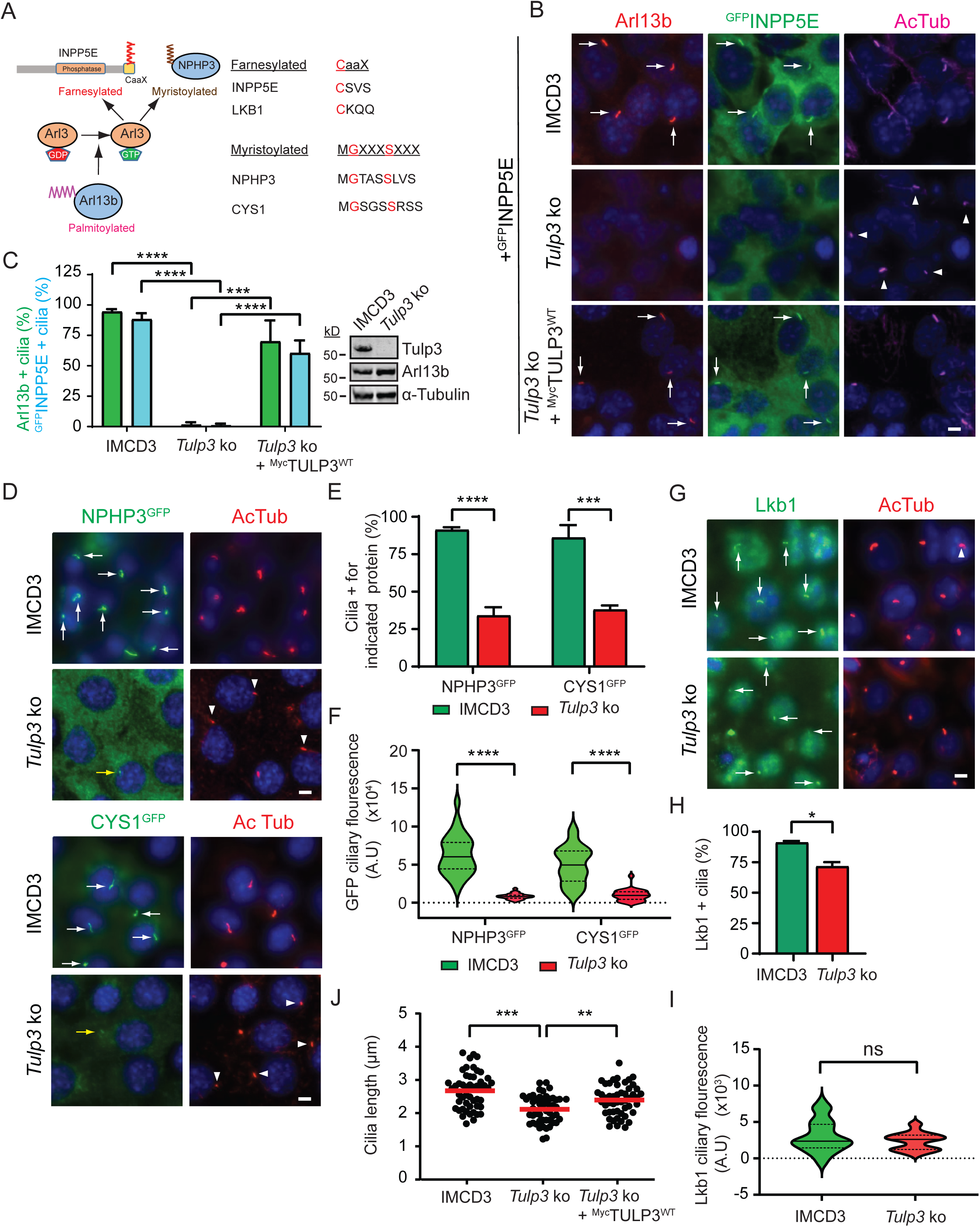
Tulp3 determines ciliary trafficking of Arl13b dependent cargoes. **(A)** Cartoon depicting the role of Arl13b in regulating the release of farnesylated and myristoylated cargoes in the ciliary compartment via Arl3 (see text). Also shows a list of lipidated proteins that are known to localize to cilia. **(B)** Wild type and *Tulp3* ko IMCD3 cells (**Figure S1B**) stably expressing ^LAP^INPP5E ± ^Myc^TULP3 were grown till confluence and serum starved for 36 hours before fixation. Fixed cells were immunostained for endogenous Arl13b, GFP (INPP5E), acetylated tubulin (AcTub) and counterstained for DNA. **(C)** Arl13b and GFP positive cilia from (B) were counted from two experiments and >200 cilia were counted for each condition. Data represent means ± SD. Western blot shows Tulp3, Arl13b and *α*Tubulin protein levels from wild type and *Tulp3* ko cells. **(D)** Wild type or *Tulp3* ko IMCD3 cells stably expressing NPHP3^LAP^ or CYS1^LAP^ were grown till confluence and serum starved for 36 hours before fixation. Fixed cells were immunostained for GFP along with acetylated tubulin (Actub) and counterstained for DNA. **(E)** GFP positive cilia from (D) were counted from two experiments, and total counted cilia are >200 for each condition. Data represent mean ± SD. **(F)** Mean GFP ciliary fluorescence intensities were measured from NPHP3^LAP^ or Cys1^LAP^ expressing wild type or *Tulp3* ko IMCD3 cells and shown as a violin plot. >30 cilia per condition were measured. **(G)** Wild type or *Tulp3* ko IMCD3 cells were grown till confluence and serum starved for 36 hours before fixation. Fixed cells were immunostained for Lkb1 along with acetylated tubulin (Actub) and counterstained for DNA. **(H)** Lkb1 positive cilia from (G) were counted from two experiments, and total counted cilia are >200 for each condition. Data represent mean ± SD. **(I)** Mean Lkb1 ciliary fluorescence intensities were measured from wild type or *Tulp3* ko IMCD3 cells. and shown as a violin plot. >30 cilia per condition were measured. **(J)** Cilia lengths were measured from indicated cells as described in (B) by acetylated tubulin immunostaining (50 cilia per condition). Scale, 5 μm. ****, p<0.0001; ***, p<0.001; **, p<0.01; *, p<0.05; Arrows indicate cilia positive for the indicated proteins while arrow heads indicate negative cilia. Yellow arrows point to cilia with low intensity of fluorescence. See also **Figure S3**.

To directly test effects from ARL13B on INPP5E localization in cilia, we stably expressed N term- or C term-GFP-Stag (LAP) tagged ARL13B in immortalized *Arl13b^hnn^* MEFs (**Figure S3**) (Larkins et al., 2011). *Arl13b^hnn^* embryos completely lack Arl13b (Caspary et al., 2007) and do not have ^HA^INPP5E in cilia upon stable overexpression. Both N- or C-tagged ARL13B fusions trafficked to cilia in *Arl13b^hnn^* background. ^HA^INPP5E levels in cilia were also restored by either fusion, suggesting rescue from their stable expression (**Figure S3**).

### Tulp3 is required for ciliary localization of lipidated proteins in murine kidneys

Embryonic cko of *Arl13b* (Li et al., 2016; Seixas et al., 2016) and potential Arl13b regulated cargoes, *Inpp5e* (Hakim et al., 2016), *Nphp3* (Bergmann et al., 2008) and *Cys1* (Hou et al., 2002) cause mild cystic disease, phenocopying *Tulp3* loss at similar timelines (Hwang et al., 2019). Therefore, we tested the role of Tulp3 in ciliary trafficking of lipidated cargoes *in vivo* in the mouse kidney. *Tulp3* ko mice are embryonic lethal and die by age E13.5 (Norman et al., 2009). Therefore, we generated conditional *Tulp3* knockouts using collecting duct-specific *HoxB7-Cre* (*HoxB7-Cre; Tulp3^f/f^)* and nephron-specific *Ksp-Cre* (*Ksp-Cre; Tulp3^f/f^)* (Shao et al., 2002; Yu et al., 2002). These mice develop progressive cystogenesis but survive till postnatal day 30 (P30) (Hwang et al., 2019). We previously showed that Arl13b and Inpp5e localized to collecting duct cilia and localization was reduced in both *Hoxb7-Cre; Tulp3^f/f^* and *Ksp-Cre; Tulp3^f/f^* collecting duct cilia by P5 (Hwang et al., 2019). Nphp3 localizes to the proximal inversin zone in cilia (Bennett et al., 2020) but localization in kidney collecting ducts is unknown. Lkb1 has been reported to localize to kidney collecting duct cilia (Viau et al., 2018). We found that both Nphp3 and Lkb1 were localized to collecting duct cilia in control mice (**Figure 3A-B**). However, both *Hoxb7-Cre; Tulp3^f/f^* and *Ksp-Cre; Tulp3^f/f^* kidney collecting ducts had reduced ciliary localization of Nphp3 by P24, whereas Lkb1 was unaffected (and **Figure 3A-B**).

**Figure 3.**
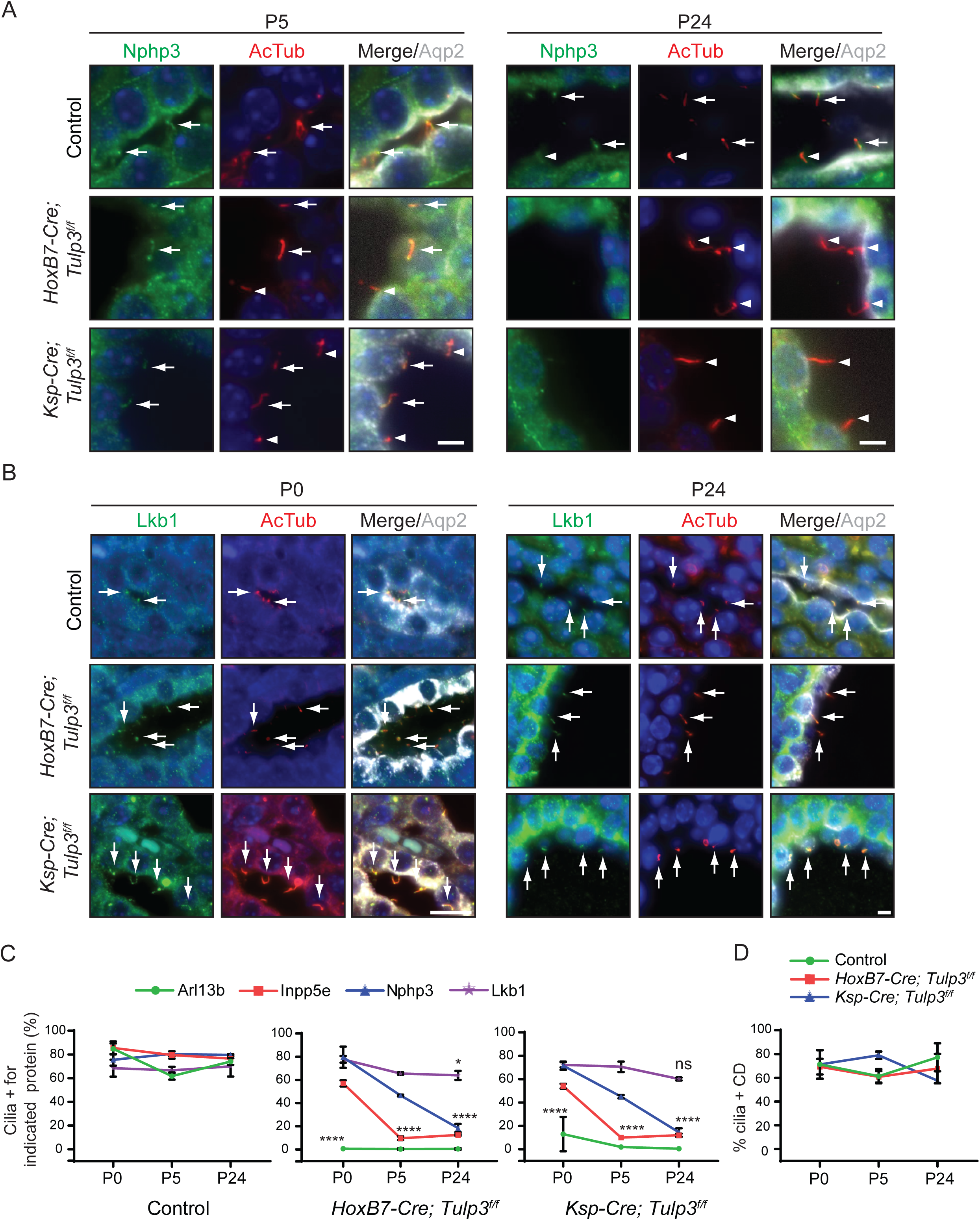
Tulp3 is required for trafficking of Arl13b-dependent lipidated cargoes to mouse kidney epithelial cilia. **(A-B)** Kidney sagittal sections from either control, *HoxB7-Cre*; *Tulp3*^f/f^ or *Ksp-Cre*; *Tulp3*^f/f^ animals at postnatal days 0, 5 and 24 were immunostained for Nphp3 **(A)** or Lkb1 **(B)**, acetylated tubulin (AcTub) and Aquaporin 2 and counterstained for DNA. **(C-D)** Percent positive cilia for the indicated proteins determined by immunostaining as in **(A-B)** were counted at postnatal days 0, 5 and 24 from 2-4 mice each genotype. Cilia counted by AcTub staining. >200 cilia were counted per mouse per genotype. Data represent mean ± SD. Scale, 5 μm. ****, p<0.0001; **, p<0.01; *, p<0.05; ns, not significant. Arrows indicate cilia positive for the indicated proteins while arrow heads indicate negative cilia.

Importantly, we observed a significant difference in the kinetics of loss of these proteins from *Tulp3* cko kidney cilia (**Figure 3C**). Arl13b is almost completely lost by P0, Inpp5e by P5 and Nphp3 by P24. This is consistent with our cell line data where Arl13b and INPP5E were the most significantly affected while NPHP3 and CYS1 were comparatively less affected (**Figure 2**). The extent of ciliogenesis was unaffected in *Tulp3* cko collecting ducts as reported before (**Figure 3D**) (Hwang et al., 2019). Overall, our data using cell lines and murine kidneys strongly suggest that Tulp3 acts as a master regulator for trafficking of different classes of Arl13b regulated lipidated proteins to cilia.

### TULP3 traffics lipidated cargoes to cilia in a predominantly PI(4,5)P_2_-independent but IFT-A-dependent manner

We previously demonstrated that both PI(4,5)P_2_ binding and IFT-A binding properties of TULP3 are required for its role in ciliary trafficking of GPCRs (Badgandi et al., 2017; Mukhopadhyay et al., 2010). We asked if PI(4,5)P_2_ and IFT-A binding properties of TULP3 are also required for trafficking Arl13b and ^HA^INPP5E to cilia. Two residues K268 and R270 in the C-terminal Tubby domain of TULP3 are required for interaction with PI(4,5)P_2_ (Mukhopadhyay et al., 2010; Santagata et al., 2001), whereas conserved residues in an N-terminal *α*-helical region of TULP3 are important for its interaction with IFT-A complex (Mukhopadhyay et al., 2010). We stably expressed mutant forms of ^LAP^TULP3 that were deficient in either PI(4,5)P_2_ binding (K268A; R270A designated as TULP3^KR^) (Mukhopadhyay et al., 2010; Santagata et al., 2001) or IFT-A binding (TULP3^mut12^) (Mukhopadhyay et al., 2010) or both PI(4,5)P_2_ and IFT-A binding (Tulp3^KR/mut12^) in *Tulp3* ko IMCD3 cells (**Figure 4A**). We observed that expression of *TULP3^mut12^* or *TULP3^KR/mut12^* did not rescue ciliary trafficking of either Arl13b or ^HA^INPP5E indicating that Tup3-IFT-A binding is required for ciliary trafficking of lipidated cargoes (**Figure 4B-D**). Interestingly, expression of *TULP3^KR^* efficiently rescued ciliary trafficking of both Arl13b and ^HA^INPP5E (**Figure 4B-D**), indicating that TULP3-PI(4,5)P_2_ binding is dispensable for lipidated cargo trafficking by TULP3. However, the ciliary intensities of both Arl13b and ^HA^INPP5E were reduced in cells rescued by TULP3^KR^, indicating a partial rescue (**Figure 4D**). In contrast, expression of *TULP3^KR^* did not rescue ciliary trafficking of Gpr161, a GPCR cargo of TULP3, similar to expression of *TULP3^mut12^* or *TULP3^KR/mut12^* (**Figure 4B-D**), confirming that both PI(4,5)P_2_ and IFT-A binding by TULP3 are required for ciliary trafficking of Gpr161 (Mukhopadhyay et al., 2010). Expression of *TULP3^KR^* also partially restored ciliary length in *Tulp3* ko cells compared to expression of either TULP3^mut12^ or TULP3^KR/mut12^ (**Figure 4E**). Recently, Han *et. al.* showed that PI(4,5)P_2_ binding property of TULP3 is not required for ciliary trafficking of ARL13B and GPR161 (Han et al., 2019). Our results using stably expressed *TULP3* WT and mutants, rather than transient expression by transfecting variants in the previous study (Han et al., 2019), suggest that TULP3 traffics Gpr161 in a PI(4,5)P_2_ dependent manner and lipidated cargoes in a predominantly PI(4,5)P_2_ independent manner.

**Figure 4.**
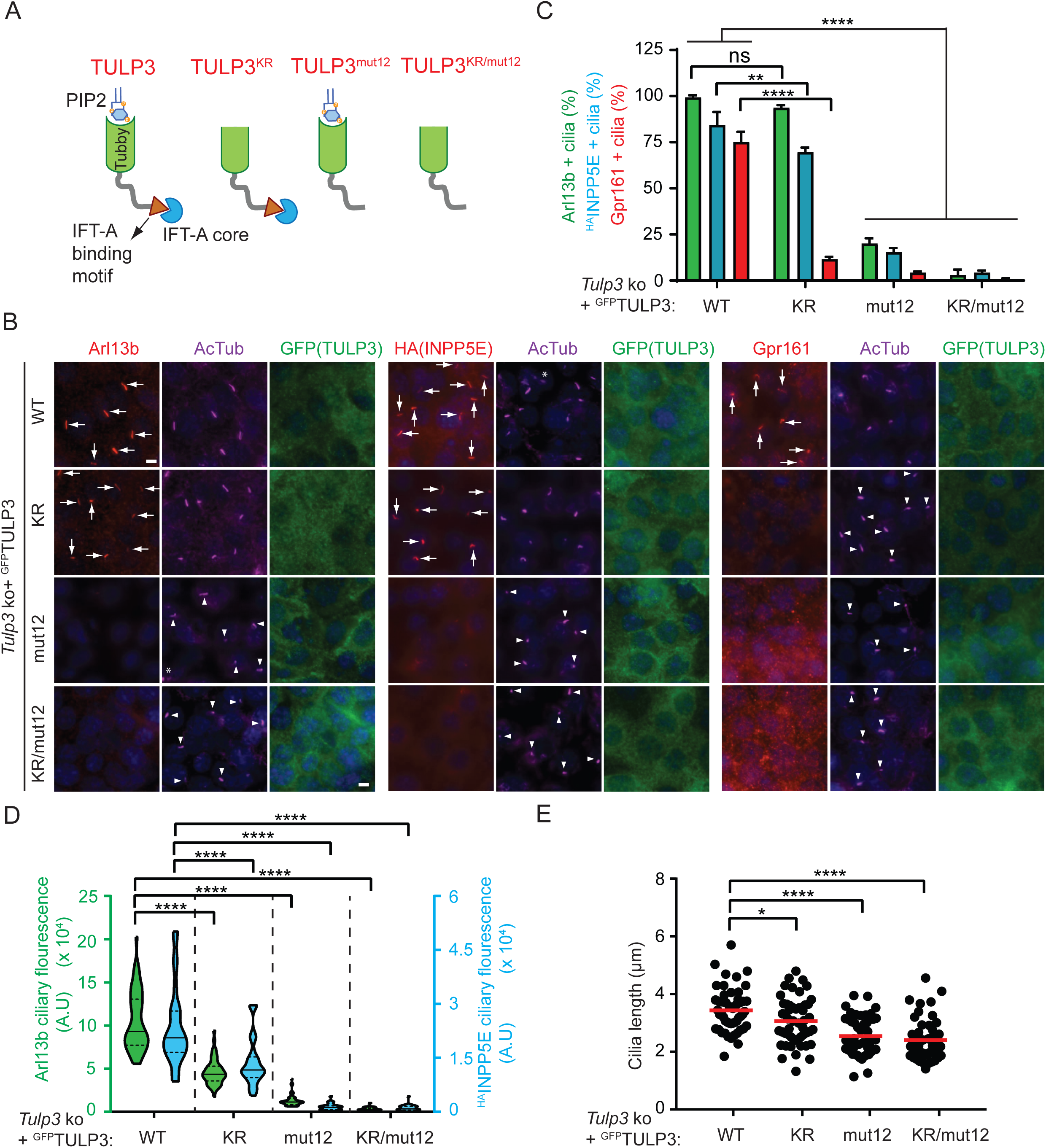
Tulp3 traffics Arl13b-dependent lipidated cargoes in a PI(4,5)P_2_ independent manner unlike transmembrane GPCR cargoes. **(A)** Schematic showing the nature of different mutants used in (B). **(B)** Wildtype or *Tulp3* ko IMCD3 line stably expressing ^HA^INPP5E and indicated ^LAP^TULP3 variants were grown to confluence and serum starved for 36 hours before fixation. Fixed cells were immunostained for endogenous Arl13b, HA (INPP5E) or endogenous Gpr161 along with GFP (TULP3) and acetylated tubulin (AcTub) and counterstained for DNA. **(C)** Arl13b/ HA/Gpr161 and GFP positive cilia were counted from two experiments, and total counted cells are >200 for each condition. Data represent mean ± SD. **(D)** Violin plot shows mean ciliary intensities of Arl13b and HA from different cell types used in (B) (50 cilia each for Arl13b and 30 cilia each for ^HA^Inpp5e). **(E)** Cilia lengths were measured from different cell types used in (A) by acetylated tubulin immunostaining (50 cilia each). Scale, 5 μm. ****, p<0.0001; **, p<0.01; *, p<0.05; ns, not significant. Arrows indicate cilia positive for the indicated proteins while arrowheads indicate negative cilia.

### Proximity biotinylation assays determine ARL13B-TULP3 interactions

As Tulp3 is required for ciliary trafficking of Arl13b, we tested if there is a physical association between the two proteins. We showed earlier that the ciliary localization sequence (CLS) fusions of transmembrane cargoes with CD8*α* are in proximity to TULP3 using proximity biotinylation or cross linking *in vivo*. (Badgandi et al., 2017) Similarly, we co-expressed ARL13B or CLSs’ fused with a promiscuous biotin ligase BirA* (BirA mutant R118G (Roux et al., 2012)) and ^LAP^TULP3 in TRex-293 cells (LAP, EGFP-TEV-Stag-X; **Figure 5A**). To remove background resulting from non-specific biotinylation of GFP, we performed tandem affinity purifications of ^LAP^TULP3. We observed that TULP3 was biotinylated by BirA*-ARL13B in comparison to a CD8*α* control without any CLS (CD8 Linker; **Figure 5B**). The biotinylation of TULP3 in presence of ARL13B was at similar levels to that in presence of the fibrocystin CLS fused with CD8*α*, one of the transmembrane cargoes of TULP3 that we identified earlier (Badgandi et al., 2017). TULP3 has a conserved C-terminal tubby domain and an N-terminal region that contains IFT-A binding motif. The C-terminal tubby domain of TULP3, but not the N-terminus, has been shown to be in proximity to its transmembrane cargoes (Badgandi et al., 2017). Similarly, we observed that the C-terminal tubby domain of TULP3, but not the N terminal region, was biotinylated by ARL13B (**Figure 5B**), indicating that ARL13B, like GPCR cargoes, interacts with the tubby domain.

**Figure 5:**
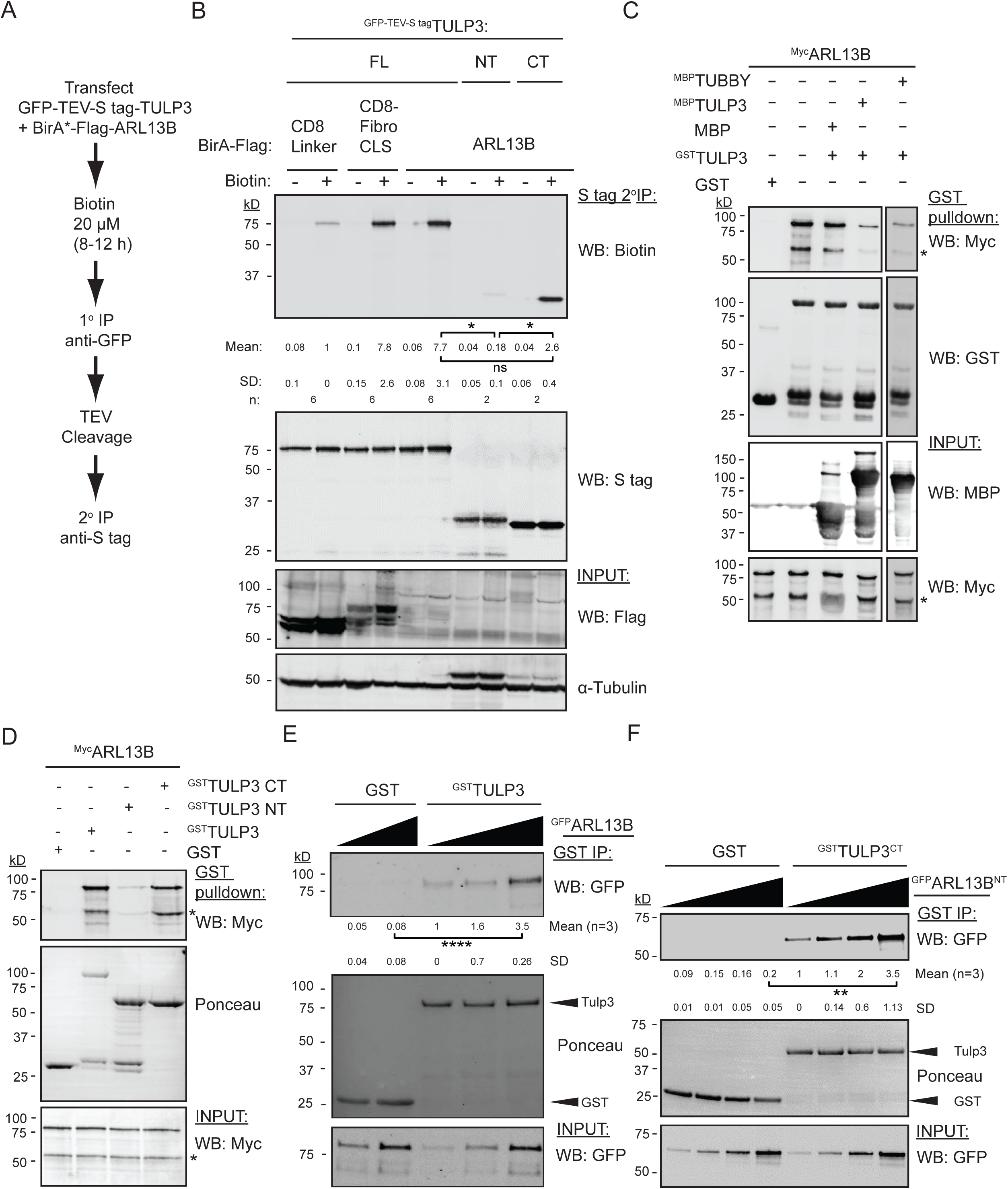
Tubby domain of TULP3 directly interacts with ARL13B. **(A)** Flowchart showing steps involved in proximity biotinylation assay in cells. **(B)** T-REx 293 cells were co-transfected with GFP-TEV-S-tagged TULP3 full-length (FL) or N-terminus (NT) or C-terminus (CT) along with BirA*-Flag tagged ARL13B and processed as shown in (A). CD8 inker is a negative control and CD8-Fibrocystin (Fibro) CLS is a positive control (Badgandi et al., 2017). Final S-tag bead eluates were processed for immunoblotting with IRdye-680 tagged streptavidin to detect biotinylation. S-tag antibody blot shows efficiency of pull down. Inputs were blotted with the indicated antibodies to detect expression. *α*Tubulin is used as a loading control. Mean ± SD values indicate Biotin/S-tag ratios normalized to CD8 linker control. **(C)** GST or ^GST^TULP3 bound beads were incubated with *In vitro* translated (IVT) ^Myc^ARL13B alone or in the presence of excess of MBP or ^MBP^TULP3 or ^MBP^Tubby protein in Lap100N at room temperature for 1 hour. Beads were washed, eluted and immunoblotted with the indicated antibodies. MBP, maltose binding protein. **(D)** *In vitro* translated ^GFP^ARL13B was incubated with GST, ^GST^TULP3 full length, ^GST^TULP3 NT or ^GST^TULP3 CT in LAP100N buffer at room temperature for one hour. Beads were processed for immunoblotting with the indicated antibodies. **(E)** Bacterially purified GST or ^GST^TULP3 protein bound to glutathione sepharose beads were incubated with insect cell purified GFP tagged ARL13B in LAP150N buffer at 4°C for 1 h. Beads were washed and immunoblotted with GFP antibody to show direct *in vitro* interaction. Ponceau staining was performed to show GST or ^GST^TULP3 on beads. Input shows flowthrough immunoblotted with GFP antibody to show the presence of ^GFP^ARL13B in the reaction. **(F)** Same as in (E), but tubby domain of TULP3 (^GST^TULP3^CT^) and insect purified GFP tagged N-terminus of ARL13B (1-298) (See Figure 8 schematic) used for *in vitro* binding reactions. Number of experiments performed denoted by “n”. Asterix in immunoblots in (C) and (D) refer to nonspecific band in anti-MYC blot in IVT reactions. ****, p<0.0001; **, p<0.01; *, p<0.05; ns, not significant.

### The Tubby domain of TULP3 binds directly to ARL13B

TULP3 traffics ARL13B in a predominantly PI(4,5)P_2_ independent manner unlike its GPCR cargoes. Therefore, we speculated that TULP3 binds to ARL13B efficiently compared to its GPCR cargoes. Further, TULP3 cargoes that are known to date are integral membrane proteins and we previously could not confirm direct interaction between these cargoes and TULP3. As ARL13B is a membrane associated protein, we tested if ARL13B interacts directly with TULP3 *in vitro*. We prepared pure ARL13B protein by *in vitro* translation (IVT) using TNT-SP6 wheat germ extract. *In vitro* translated ^Myc^ARL13B interacted with bacterially purified ^GST^TULP3 (**Figure 5C**). Excess ^MBP^TULP3 or ^MBP^Tubby, but not MBP alone, could compete out ^Myc^ARL13B binding to ^GST^TULP3 (**Figure 5C**), suggesting that conserved regions in Tubby family proteins might be involved in interaction with ARL13B. Furthermore, using bacterially purified GST tagged TULP3 N and C terminal truncations, we confirmed that the C-terminal tubby domain of TULP3 interacts with *in vitro* translated ^Myc^ARL13B (**Figure 5D**). Thus, the C-terminus tubby domain of TULP3 directly binds to ARL13B.

Our attempts to purify ARL13B from bacterial cells were not successful as we always saw multiple degradation products in our protein preparations. Thus, we purified full length and N-term fragment (1-298 aa; ARL13B^NT^) of ARL13B from Sf9 insect cells using the FlexiBac system (Lemaitre et al., 2019). The Sf9 cell system also allows purification of proteins with posttranslational modifications such as palmitoylation (Mouillac et al., 1992; Ng et al., 1994). The full length ARL13B directly bound with ^GST^TULP3 *in vitro*. The bound fraction increased proportionally to the amount of ARL13B in the reactions (**Figure 5E**). Furthermore, such binding was also seen between the tubby domain of TULP3 (^GST^TULP3^CT^) and ARL13B^NT^ (**Figure 5F**), with bound fractions increasing proportionally to the levels of ARL13B^NT^ in the reactions. Thus, TULP3 directly binds through its tubby domain to ARL13B.

### A conserved lysine in the tubby domain of TULP3 is crucial for cargo interactions

To identify potential cargo binding residues on TULP3 we performed proximity based biotinylation using BirA* tagged GPCR CLSs followed by mass spectrometry. We identified multiple lysine residues in the tubby domain of TULP3 that were in proximity to the GPCRs Gpr161 and MCHR1 in comparison to corresponding mutated CLSs that are non-ciliary (**Figure S4A**) (Badgandi et al., 2017). One of these lysine residues, K389 in human TULP3 corresponds to K407 in mouse Tulp3 (**Figure S4B**) and was also identified as a *Tulp3^K407I^* allele in mice exhibiting perinatal lethality and kidney cystogenesis during later embryogenesis (Legue and Liem, 2019). This allele is hypomorphic to *Tulp3* null that is embryonic lethal at E14.5 (Norman et al., 2009), but it phenocopies *Tulp3* cko in the embryonic kidney that causes postnatal cystogenesis (Hwang et al., 2019). To test if this residue in TULP3 is required for cargo binding, we performed cell-based proximity biotinylation. Interestingly, we noticed that ^LAP^TULP3^K389I^ had significantly reduced proximity to ARL13B compared to ^LAP^TULP3^WT^ (**Figure 6A**). Furthermore, we observed that ^GST^TULP3^K389I^ had reduced direct binding with insect cell purified ARL13^NT^ (1-298 aa) *in vitro* compared to ^GST^TULP3 (**Figure 6B**). Thus, lysine 389 in TULP3 is required for direct interactions with ARL13B.

**Figure 6:**
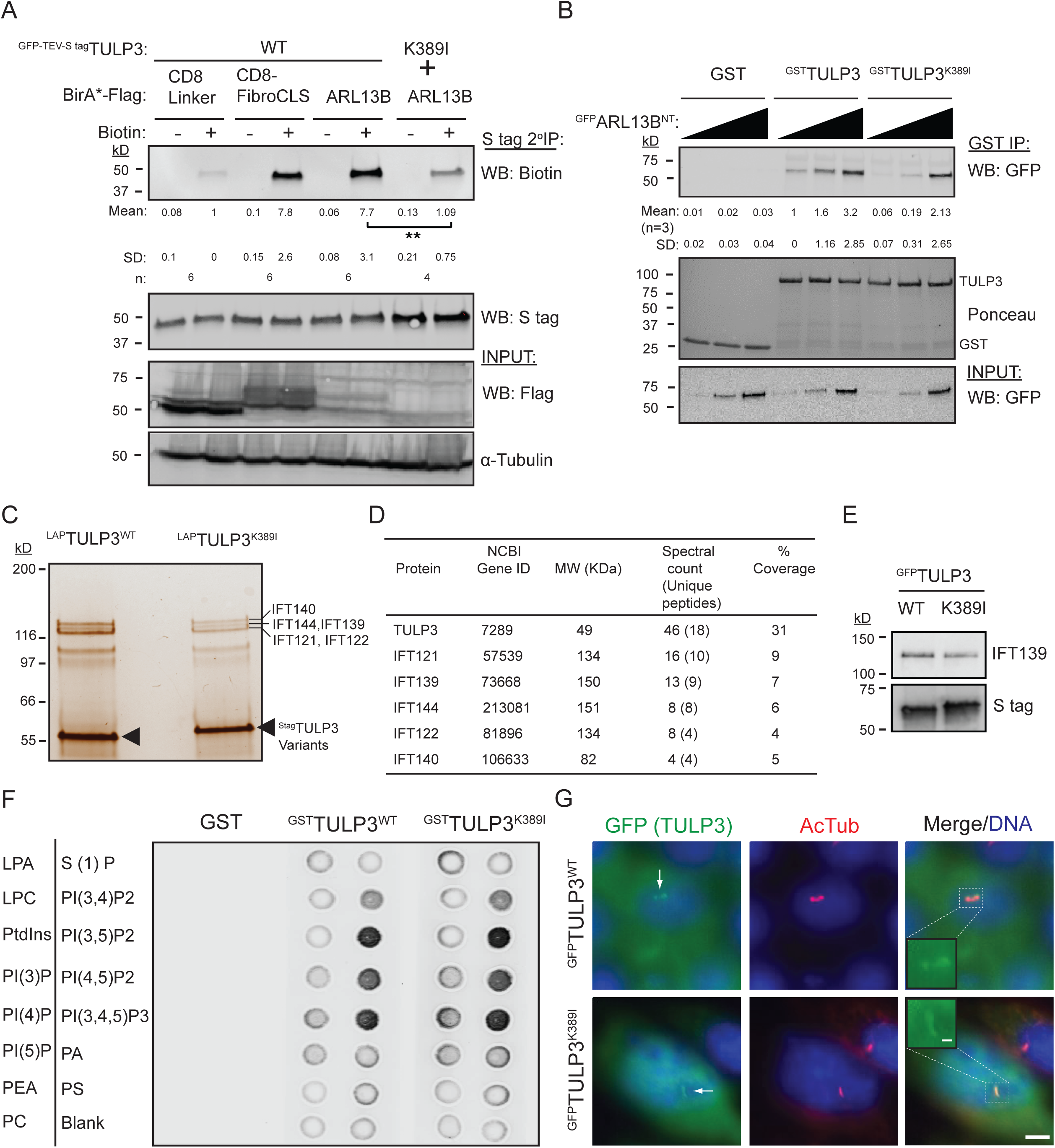
Lysine K389 in TULP3 is required for interaction with its cargoes. **(A)** T-Rex 293 cells were co-transfected with GFP-TEV-S-tagged TULP3 wild type (WT) or K389I mutant along with BriA*-Flag tagged ARL13B and processed as shown in 5A-B. “n” indicates number of experiments performed. Data represent mean ± SD. **(B)** Increasing amounts of insect cell purified ^GFP^ARL13B (1-298) (NT) protein was incubated with either GST, ^GST^TULP3 or ^GST^TULP3^K389I^ as in 5E-F. “n” indicates number of experiments performed. Data represent mean ± SD. **(C)** Tandem affinity purification of RPE cells stably expressing LAP tagged wild type TULP3 or K389I mutant was performed, and the silver stain shows the presence of multiple IFT-A components in the pull downs. **(D)** Samples from (C) were subjected to mass spectrometry to identify the proteins in the pulldown complexes. Table shows the IFT-A components identified along with respective peptide counts and percentage of coverage. **(E)** Samples from (C) were subjected to immunoblotting with anti-IFT139 (THM1) and S-tag antibody. **(F)** Equal amounts of GST, ^GST^TULP3 or ^GST^TULP3^K389I^ protein was added to lipid strips to test for lipid binding properties of the proteins. **(G)** IMCD3 cells stably expressing ^LAP^TULP3^WT^ or ^LAP^TULP3^K389I^ were serum starved for 36 hours before fixing and immunostained for GFP, Acetylated tubulin and DNA. Insets hsow TULP3 localization in cilia. Scale, 5 μm. Inset scale, 1 μm. **, p<0.01. See also **Figure S4**.

TULP3 has multiple other cellular roles including binding to the IFT-A core through the N-terminus, binding to phosphoinositides including PI(4,5)P_2_ through the tubby domain, and ciliary localization (Mukhopadhyay et al., 2010). We identified most IFT-A complex proteins among interacting partners of stably expressed LAP-tagged TULP3^K389I^ in RPE-hTERT cells using tandem affinity purification and mass spectrometry (**Figure 6C-6D**) and by immunoblotting for IFT139 (**Figure 6E**). We found that recombinant ^GST^TULP3^K389I^ also bound to select phosphoinositides including PI(4,5)P_2_ on PIP blots similar to ^GST^TULP3^WT^ as reported before (**Figure 6F**) (Mukhopadhyay et al., 2010). Finally, ^LAP^TULP3^K389I^ was localized to primary cilia upon stable expression in IMCD3 cells by immunofluorescence (**Figure 6G**) as for the wildtype protein (Mukhopadhyay et al., 2010). Thus, TULP3^K389I^ had diminished cargo binding, whereas phosphoinositide binding, IFT-A binding and ciliary localization remained unaffected.

### Lysine 389 in TULP3 is required for ciliary trafficking of its cargoes

We next tested if the K389 residue in TULP3 is important for ciliary cargo trafficking. We stably expressed *TULP3^WT^* or *TULP3^K389I^* in *Tulp3* ko IMCD3 cells and observed that WT, but not K389I, could rescue ciliary localization of Arl13b, ^HA^INPP5E and Gpr161 (**Figure 7A-C**). The ciliary length in *TULP3^K389I^* expressing cells showed a small reduction in length compared to *TULP3^WT^* expressing cells (**Figure 7D**). Furthermore, endogenous ciliary pools of both Arl13b and Gpr161 were abrogated in MEFs from *Tulp3^K407I^* mice (**Figure 7E-F**). Thus, the lysine 389 in TULP3 was critical for trafficking both GPCR and lipidated cargoes to cilia suggesting that a common interface in the tubby domain of TULP3 mediated trafficking of both transmembrane and membrane associated cargoes.

**Figure 7:**
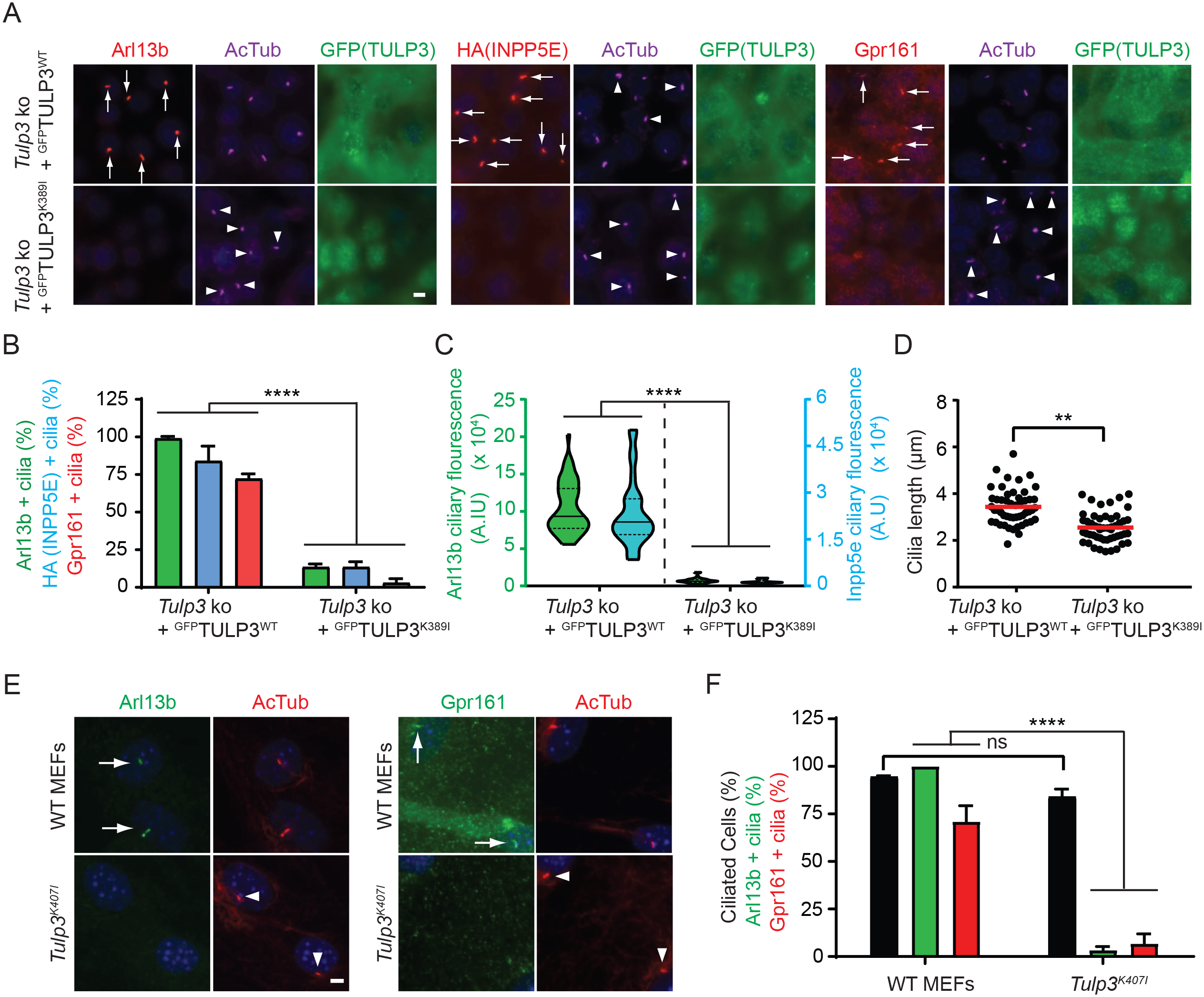
Lysine 389 in TULP3 is required for ciliary cargo trafficking. **(A)** Wildtype or *Tulp3* ko IMCD3 line stably expressing ^HA^INPP5E and LAP-tagged wild type or K389I mutant of *TULP3* were grown till confluence. Cells were serum starved for 36 h before fixation and then immunostained for Arl13b or HA or Gpr161 along with GFP, acetylated tubulin (AcTub) and counterstained for DNA. **(B)** Arl13b/HA/Gpr161 in GFP(TULP3) positive cilia were counted from two experiments, and total counted cells are >200 for each condition. Data represent mean ± SD. **(C)** Violin plot shows mean ciliary intensities of Arl13b and HA (50 cilia each for Arl13b and 30 cilia each for ^HA^INPP5E). **(D)** Cilia lengths were measured from by immunostaining for acetylated tubulin (>50 cilia per condition). **(E)** MEFs from wild type or *Tulp3^K407I^* mice were serum starved for 24 h before fixation and immunostained for either Arl13b or Gpr161 along with acetylated tubulin (AcTub) and counterstained for DNA. **(F)** Arl13b or Gpr161 positive cilia were counted from two experiments, and total counted cells are >200 for each condition. Data represent mean ± SD. Scale, 5 μm. ****, p<0.0001; **, p<0.01; ns, not significant. Arrows indicate cilia positive for the indicated proteins while arrowheads indicate negative cilia.

### The N-terminus of ARL13B mediates TULP3 binding

Our earlier studies showed that for almost all TULP3 dependent cargoes, the ciliary targeting motifs are short sequence in the respective proteins. These sequences are in proximity to TULP3 and are both necessary and sufficient for ciliary trafficking by TULP3 (Badgandi et al., 2017). To dissect ciliary targeting of ARL13B by TULP3, we aimed at identifying ARL13B regions involved in TULP3 binding. ARL13B has a palmitoylation motif and GTPase domain followed by a coiled-coiled (CC) domain in its N-terminus along with RVxP motif and Proline Rich (PR) region in its C-terminus (**Figure 8A**). We tested various *in vitro* translated truncations of ^Myc^ARL13B for their capacity to bind with ^GST^TULP3 and observed that the N-terminus of ARL13B corresponding to the GTPase and coiled coil domains (D2, 1-244 aa) binds to TULP3 stronger than the C-terminus alone (D5, 245-428 aa) (**Figure 8A-B**). As mentioned before, the insect cell purified N-terminus (residues 1-298) of ^GFP^ARL13B also binds to the bacterially purified C-terminal tubby domain of ^GST^TULP3 *in vitro* (**Figure 5F**). Therefore, the N-terminus of ARL13B mediates direct TULP3 binding.

**Figure 8:**
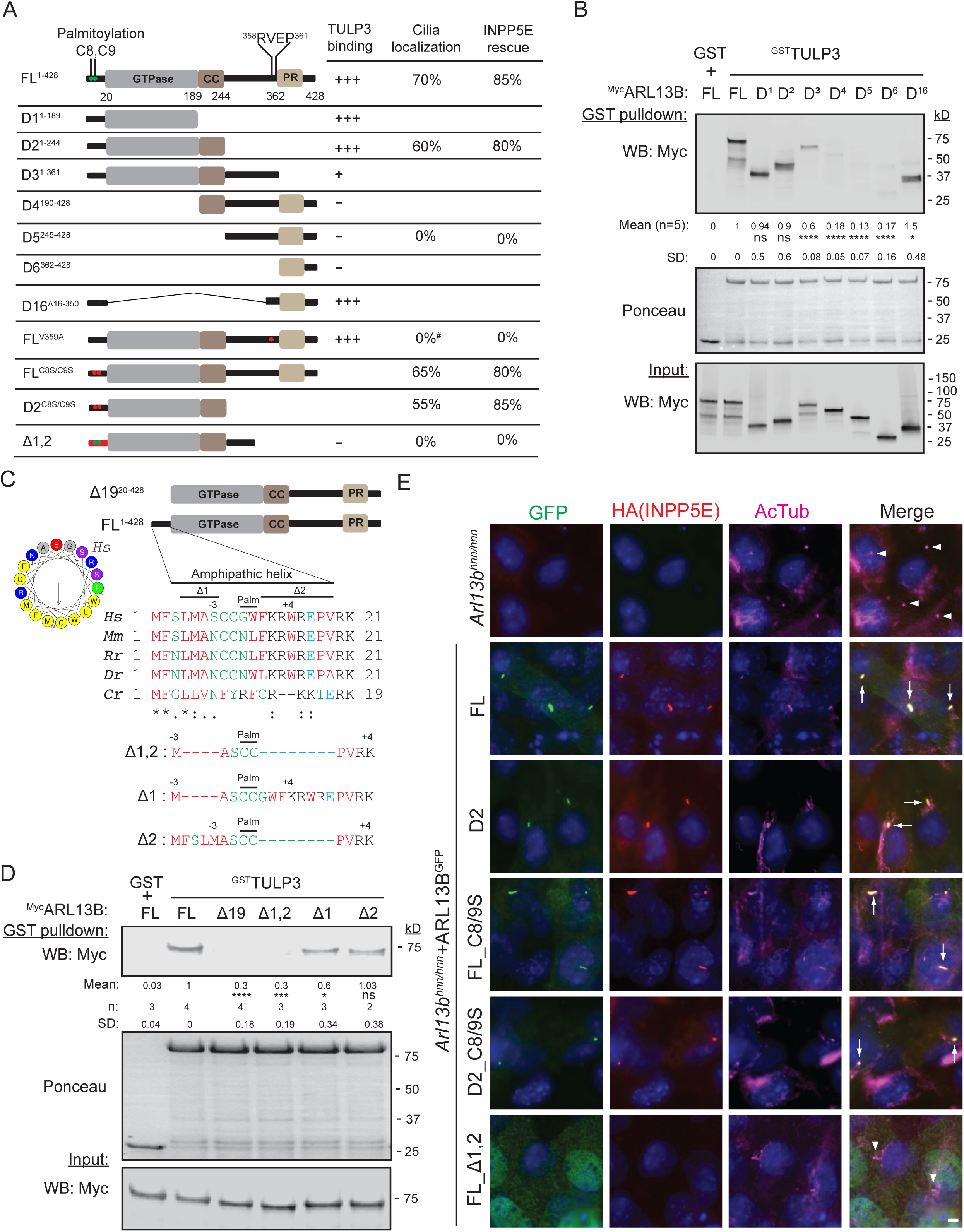
Arl13b domains required for Tulp3 interaction and ciliary localization. **(A)** Schematic representation of different ARL13B truncations tested for *in vitro* TULP3 binding (“TULP3 binding”) or stably expressed in immortalized *Arl13b^hnn^* MEFs along with HA-tagged INPP5E to test for ciliary trafficking (“cilia localization”) and rescue of ARL13B function (“INPP5E rescue”), respectively. ∼30-100 cilia counted for cilia localization and HA-tagged INPP5E positive cilia. *Arl13b^hnn^* MEFs had no HA-tagged INPP5E in cilia. ^#^, data from localization of endogenous mutant Arl13b^V358A^ and Inpp5e in *Arl13b^V358A/V358A^* MEFs (Figure S5B) (Gigante et al., 2020). **(B)** Bacterially purified GST or ^GST^TULP3 protein bound to glutathione Sepharose beads was incubated with *in vitro* translated (IVT) Myc tagged truncations of ARL13B in Lap150N buffer at room temperature for 1 h. Beads were washed, eluted and immunoblotted with Myc antibody to show interaction. Ponceau stain shows GST or ^GST^TULP3 on beads. Input shows flowthrough immunoblotted with GFP antibody to show the presence of ^GFP^ARL13B in the reaction. Conserved palmitoylation motif residues with respect to the palmitoylated Cysteine are shown in each deletion (Ren et al., 2008; Weng et al., 2017) **(C)** Alignment of N-terminus region of ARL13B preceding the GTPase domain and mutants tested for TULP3 binding and for ciliary localization. ARL13B protein IDs: Hs, homo sapiens NP_001167621.1; Mm, *Mus musculus* NP_080853.3; Rn, *Rattus norvegicus* NP_001100571.1; Dr, *Danio rerio* NP_775379.1; Cr, *Chlamydomonas reinhardtii* A8INQ0.1. Amphipathic helix for the human protein drawn using HeliQuest (Gautier et al., 2008). Arrow indicates hydrophobic moment. **(D)** Bacterially purified GST or ^GST^TULP3 protein bound to glutathione Sepharose beads was tested for binding with *in vitro* translated Myc tagged full length ARL13B and indicated mutants as in (B). **(E)** *Arl13b^hnn^* cells stably expressing ^HA^INPP5E with indicated C-LAP tagged ARL13B fusions were serum starved for 48 h before fixing and immunostained for GFP, HA (INPP5E), acetylated tubulin (AcTub) and counterstained for DNA. Quantification in (A) and Figure S5A. Scale, 5 μm. ****, p<0.0001; ***, p<0.001; *, p<0.05; ns, not significant. Arrows indicate cilia positive for the indicated proteins while arrowheads indicate negative cilia. See also **Figure S5**.

### The C-term RVxP motif of ARL13B requires membrane anchoring for trafficking to cilia in a TULP3 independent manner

Previous studies reported the RVxP motif (sequence R^357^VEP) in the C-terminus of mouse Arl13b (corresponding to 358-361 R^358^VEP of human ARL13B) as a potential ciliary localization signal (Mariani et al., 2016). As reported before (Gigante et al., 2020), we did not see the RVxP mutant protein in cilia in MEFs with knock-in at the endogenous RVxP locus (*Arl13b^V358A/V358A^*) (**Figure S5A**). By cell-based proximity biotinylation assay, we noted that ARL13B^V359A^ interacted with TULP3 efficiently when compared with the wild type ARL13B (**Figure S5B**). Similarly, *in vitro* binding between ^GST^TULP3 and *in vitro* translated ^Myc^ARL13B^V359A^ was similar compared to wild type ^Myc^ARL13B (**Figure S5C**). Thus, ARL13B RVxP mutant failed to traffic to cilia despite efficient TULP3 binding.

We next tested additional requirements for the RVxP motif to traffic to cilia. A *β*1 Integrin-HaloTag-ARL13B–C-GFP (IA-GFP) traffics to cilia (Su et al., 2013). In this construct the C-term region of human ARL13B containing the RVxP motif (355–428 aa) is anchored to the cell surface by fusion to the transmembrane domain of human β1 integrin (Svendsen et al., 2008). By transfecting IA-GFP in *Tulp3* ko NIH 3T3 cells, we found that this construct is trafficked to cilia independent of Tulp3 (**Figure S5D**), as expected from the corresponding C-terminus domain of ARL13B (D5 and D6, 362-428) not binding to TULP3. (**Figure 8A-B**). The *β*1 Integrin-HaloTag-GFP was not by itself able to traffic to cilia (**Figure S5D**). As reported earlier (Larkins et al., 2011), the ARL13B D5 fragment fused with C-terminal LAP was not trafficked to cilia when expressed in *Arl13b^hnn^* MEFs (**Figure 8A, S5E**). Therefore, membrane anchoring is required for the RVxP motif containing C-term ARL13B fragment to traffic to cilia and is independent of TULP3.

### An N-terminal amphipathic helix in ARL13B binds to TULP3

Based on other ARF family proteins (Donaldson and Jackson, 2011; Randazzo et al., 1995), ARL13B has a lipid-sensitive clamp provided by an amphipathic helix at the N terminus preceding the GTPase domain. Thus, the first 19 residues of human/mouse ARL13B (Hori et al., 2008; Ivanova et al., 2017) and corresponding residues in the *Chlamydomonas* protein (Gotthardt et al., 2015) were not included when performing GTP binding assays and structural studies. We already had the first 19 aa in our insect cell purified ^GFP^ARL13B protein that showed binding with TULP3 (**Figure 5E-F** and **6B**). We next tested if the C-terminal region fused with the short N-terminal amphipathic helix (D16, **Figure 8A**) interacts with TULP3. We observed that this construct bound to TULP3 as efficiently as the full-length or the N-terminus (D2) *in vitro* (**Figure 8B**).

The amphipathic conserved helix flanks the palmitoylation site (**Figure 8C**). We wondered if the amphipathic helix was responsible for TULP3 binding. We first tested the helix deleted ^Myc^ARL13B^Δ19^ in our *in vitro* translated protein binding assays (**Figure 8C**). Remarkably, this protein was almost fully abrogated in its binding to ^GST^TULP3 (**Figure 8D**). Deletion of 4 conserved aa preceding the palmitoylation site (Δ1; **Figure 8C**) and but not of seven conserved aa following the palmitoylation site (Δ2; **Figure 8C**) in ^Myc^ARL13B modestly reduced binding to ^GST^TULP3 (**Figure 8D**). Finally, a double deletion mutant of these regions in the full-length protein (Δ1,2; **Figure 8C**) was almost fully abrogated in binding to ^GST^TULP3 (**Figure 8D**). Thus, the helix flanking the palmitoylation site mediates TULP3 binding.

### The amphipathic helix directs trafficking to cilia even in the absence of palmitoylation and RVxP motif

To directly test role of TULP3 binding in ciliary localization of ARL13B, we stably expressed different ARL13B fragments and mutants fused with C-terminal LAP in the *Ar13b^hnn^* background (**Figure 8E**). We chose to test expression in the *Ar13b^hnn^* background as self-association of ARL13B through its N-terminal domain has been reported using Co-IP experiments (Hori et al., 2008). We also co-expressed ^HA^INPP5E to check for restoration of ARL13B function in the *Ar13b^hnn^* background (**Figure 8E**).

We first tested the D2 fragment for trafficking to cilia in the *Arl13b^hnn^* background. Consistent with our TULP3 binding data (**Figure 8B**), we noticed ciliary trafficking of the D2 fragment (**Figure 8A, 8E**). ^HA^INPP5E levels in cilia were rescued (**Figure 8A, 8E**) and ciliary length was partially rescued compared to the *Ar13b^hnn^* MEFs (**Figure S5F**). Thus, the N-terminus of ARL13B (up to the CC domain) containing the TULP3 binding amphipathic helix traffics to cilia and is functional.

A full length palmitoylation deficient mutant upon transient expression by transfection has been reported to be only partially trafficked to cilia in the *Arl13b^hnn^* background (Mariani et al., 2016). Similar motif changes in *C. elegans*, however, does not prevent ARL13B trafficking to proximal segments of phasmid cilia, although promoting mislocalization to the cell body (Cevik et al., 2013). Upon stable expression in *Arl13b^hnn^*, we found that the ARL13B^C89S^ mutant fused with C-terminal LAP was trafficked to cilia. ^HA^INPP5E levels in cilia were rescued (**Figure 8A, 8E**) with partial rescue of ciliary length compared to the *Ar13b^hnn^* MEFs (**Figure S5F**). Thus, the full length ARL13B is localized to cilia despite lacking in palmitoylation. To rule out any role of palmitoylation in the D2 context, we tested if the palmitoylation deficient D2 fragment (D2^C8/9S^) trafficked to cilia (**Figure 8E**). We found that D2^C8,9S^ still trafficked to cilia in the *Arl13b^hnn^* background (**Figure 8E**). ^HA^INPP5E levels in cilia were rescued **Figure 8A, 8E**), but ciliary length was unchanged compared to the *Ar13b^hnn^* MEFs (**Figure S5F**). Thus, the amphipathic helix in ARL13B mediates binding to TULP3 and directs trafficking to cilia even in the absence of palmitoylation and the RVxP motif.

Finally, we generated a full length ARL13B fusion that lacked in TULP3 binding (ARL13B^Δ1, Δ2^) without disrupting the palmitoylation motif (Ren et al., 2008; Weng et al., 2017) (**Figure 8C**). This mutant was not trafficked to cilia (**Figure 8E**). Correspondingly, ^HA^INPP5E levels in cilia were not rescued (**Figure 8E**). Thus, TULP3 binding is absolutely required for trafficking of the full length ARL13B to cilia.

## Discussion

ARL13B is indisputably one the most used ciliary membrane markers since its discovery more than a decade ago (Caspary et al., 2007; Sun et al., 2004). The highly enriched ciliary pools of ARL13B seen in multiple tissues and ciliated cells contributes to the universality of its use as a ciliary marker. However, the mechanisms that traffic ARL13B to cilia are not well understood, and whether a defined region in ARL13B by itself suffices for ciliary trafficking is not known. We show that TULP3 is a master regulator for trafficking ARL13B to cilia without affecting total cellular pools. ARL13B trafficking to cilia is primarily regulated by direct binding to TULP3 and TULP3’s binding to the IFT-A core complex. An N-terminus amphipathic helix that flanks the palmitoylation site and precedes the GTPase domain mediates binding of ARL13B to TULP3 and directs trafficking to cilia even in the absence of palmitoylation and the RVxP sorting motif. Unlike trafficking of GPCRs that requires both IFT-A and PI(4,5)P_2_ binding by TULP3 (Mukhopadhyay et al., 2010), the binding between ARL13B and TULP3 is of sufficient strength to traffic ARL13B and ARL13B-dependent lipidated cargoes even without PI(4,5)P_2_ anchoring by TULP3. A conserved lysine in TULP3’s tubby domain (lysine 389 in human TULP3) mediates direct ARL13B binding and determines trafficking of both lipidated and transmembrane cargoes. Overall, our results suggest that TULP3 binding to short sequences in diverse cargoes is mediated by a shared region in the tubby domain.

### TULP3 enriches ARL13B-dependent lipidated proteins in cilia

We demonstrate that Arl3-dependent farnesylated (Inpp5e) and myristoylated (Nphp3, Cys1) proteins are regulated by Tulp for ciliary localization. The farnesylated protein Lkb1, which is Arl3 independent for trafficking to cilia, is not Tulp3 regulated. Inpp5e loss in cilia of *Tulp3* cko kidney collecting duct epithelia accompanies cystogenesis and follows Arl13b depletion. Nphp3 and Cys1 are reduced in ciliary intensity but retain weak levels in cilia in *Tulp3* ko cells. Nphp3 is also slow to be depleted from *Tulp3* cko kidney collecting duct epithelial cilia. The effectiveness for complete depletion of Inpp5e in cilia in *Tulp3* ko might be related to direct binding between Arl13b and Inpp5e and requirement of such binding for effective ciliary retention of Inpp5e (Humbert et al., 2012; Qiu et al., 2021). Nphp3 is concentrated in the proximal ciliary inversin compartment by binding to Nek8 and Anks6 that are required downstream of Inversin for Nphp3 localization (Bennett et al., 2020). Such binding might promote some retention of Nphp3 even in the absence of Tulp3. Nonetheless, the depletion of lipidated cargoes with distinct kinetics from kidney epithelial cilia following *Tulp3* deletion strongly suggests their differential roles in renal cystogenesis from Arl13b (Li et al., 2016; Seixas et al., 2016) or Tulp3 loss during embryogenesis (Hwang et al., 2019; Legue and Liem, 2019). The ciliary levels of Arl13b are however unaffected upon *Tulp3* deletion in brain/cerebellum and neural tube (Ferent et al., 2019), likely from redundancy between Tulp3 and Tubby in trafficking Arl13b. Such redundancy is also seen in trafficking of a subset of GPCRs by tubby in brain (Badgandi et al., 2017).

### An amphipathic helix determines ciliary localization of ARL13B

The amphipathic helix region is a signature feature in the ARF family proteins (Donaldson and Jackson, 2011; Randazzo et al., 1995) that ensures that they are associated with the membrane in GTP bound form (Antonny et al., 1997). This region had been previously omitted when performing GTP binding assays and structural studies using human, mouse (Hori et al., 2008; Ivanova et al., 2017) or *Chlamydomonas* ARL13B. We infer that the amphipathic helix is required for trafficking to cilia irrespective of palmitoylation and even in presence of intact RVxP motif from the following observations. First, a full length ARL13B mutant (ARL13B^Δ1, Δ2^) that does not bind to TULP3 but retains palmitoylation and RVxP motifs is not trafficked to cilia, suggesting TULP3 binding to be absolutely required for trafficking of the full length ARL13B to cilia. Second, an N-terminal fragment of ARL13B including just the helix, GTPase and coiled coil domains (D2, 1-244 aa) and lacking the RVxP motif traffics to cilia and rescues INPP5E levels in cilia of *Arl13b* null cells. This N-terminal fragment, even upon being non-palmitoylated, traffics into cilia, suggesting that only TULP3 binding is required for ciliary trafficking of this RVxP motif lacking fragment.

### The RVxP motif cooperates with ARL13B bound TULP3-IFT-A to increase ciliary delivery

The C-terminal RVxP motif in ARL13B was initially discovered to be involved in post-Golgi trafficking of Rhodopsin (Deretic et al., 2005). We find that the ARL13B mutated in the RVxP motif (ARL13B^V359A^) retained binding to TULP3 and proximity with TULP3 *in vivo*. However, despite TULP3 binding, the mutant protein was not trafficked to cilia in the *Arl13b^V358A/V358A^* MEFs. This result places the RVxP mechanism downstream of TULP3-IFT-A core coupling. We show that the RVxP motif by itself can traffic to cilia independent of TULP3, but only if provided with a membrane anchor (as provided by the *β*1 integrin in the IA-GFP chimera). While the actual mechanism of the RVxP motif in ARL13B is not known, for secretory proteins such as Rhodopsin, interaction between this motif and ARF4 has been proposed to be important for post-Golgi trafficking (Wang et al., 2012). Despite being cytosolic, Arl13b localizes to tubular-vesicular structures of the Arf6- and Rab22a-dependent recycling pathway (Barral et al., 2012). While ARF4 is seen among ARL13B interacting proteins in human kidney epithelial cells in an unbiased proximity-dependent biotinylation screen (He et al., 2018), the relevance of ARF4 and secretory pathway components in ARL13B trafficking to cilia would require further study. Both the N-terminus ARL13B D2 fragment without RVxP motif, and the non-palmitoylated full length ARL13B partially rescued ciliary length in *Arl13b* null cells. Levels of Arl13b correspond with ciliary length by inducing membrane protrusion and axonemal extension (Lu et al., 2015). Therefore, in addition to the TULP3 binding amphipathic helix, other motifs, such as RVxP and palmitoylation site, ensure the full complement of ARL13B in cilia and determine steady state ciliary length. Differential levels of Arl13b and downstream lipidated proteins in cilia could underlie the context specific requirements of ciliary pools of Arl13b in the kidney tubules (Duldulao et al., 2009) vs the neural tube (Gigante et al., 2020).

### Role of IFT-A in ARL13B trafficking to cilia

Apart from our results showing that TULP3-IFT-A core binding is required for ARL13B trafficking to cilia, evidence from other labs also suggests an important role for IFT-A core subunits in ARL13B ciliary localization. The IFT-A core comprises of IFT144, IFT122 and IFT140, whereas IFT43, IFT139 and IFT121 are peripheral subunits (Behal et al., 2012; Hirano et al., 2017; Mukhopadhyay et al., 2010). ARL13B levels in cilia are reduced in both *IFT122* (Takahara et al., 2018) and *IFT144* (Kobayashi et al., 2021) knockout cell lines. A recent proteomics study showed that the flagellar levels of multiple ARF family GTPases such as ARL13B, and other myristoylated and farnesylated proteins were reduced in an *IFT-140* null *Chlamydomonas* mutant (Picariello et al., 2019). Whether tubby family proteins (Stolc et al., 2005; VanderWaal Mills, 2011) underlie the anterograde trafficking of lipidated proteins by IFT-140 in *Chlamydomonas* is currently unclear. Other than the IFT-A core-TULP3 interactions, the IFT121 peripheral subunit is also required for trafficking of GPCRs, ARL13B and SMO (Fu et al., 2016). Such dependance is abrogated in the absence of F-actin depolymerization by cytochalasin D (Fu et al., 2016), suggesting that IFT121 delivers cargoes to cilia by coordinating with actin binding proteins in the face of an intact filamentous actin network. As SMO trafficking to cilia is TULP3-IFT-A core independent (Mukhopadhyay et al., 2010; Norman et al., 2009; Qin et al., 2011), the role of IFT121 peripheral subunit in trafficking SMO, and by extension ARL13B and GPCRs, is unrelated to TULP3-IFT-A core interactions.

### A shared tubby domain interface in cargo trafficking by TULP3

We find a site distinct from the PI(4,5)P_2_ binding site in the tubby domain mediating cargo binding by TULP3. Direct binding between the TULP3^K389I^ tubby domain mutant and ARL13B is reduced, as is proximity inside cells using biotinylation assays compared to wild type TULP3. PI(4,5)P_2_ binding by the tubby domain is unaffected in TULP3^K389I^, as is ciliary trafficking that is mediated by IFT-A core binding to TULP3 N-terminus (Mukhopadhyay et al., 2010). Strikingly, an equivalent mouse mutation in *Tulp3* (*K407I*) prevents trafficking of Arl13b and Gpr161 to cilia in cultured cells. Also, Arl13b trafficking to kidney epithelial cilia is affected in *Tulp3^K407I^* E18.5 kidneys (Legue and Liem, 2019). Our experiments showing direct binding between purified TULP3 and ARL13B and the discovery of an amphipathic helix in N-term of ARL13B mediating TULP3 binding, suggest a short sequence in coupling with TULP3 tubby domain. Similar diverse CLSs from cilia-targeted GPCRs and fibrocystin are also required in the membrane-bound context for proximity between cargoes and TULP3. As ciliary trafficking of both Arl13b and Gpr161 are disrupted in the *Tulp3^K407I^* MEFs, TULP3 binding to short sequences in either protein is likely to be mediated by this region in the tubby domain. The minimal requirement of these diverse sequences might be to generate secondary structure, such as for the amphipathic helix in ARL13B. Drugging this interaction domain could provide therapeutics in ciliary trafficking-regulated diseases such as polycystic kidney disease and peripheral obesity.

## Materials and methods

### Mouse strains

All mice were housed at the Animal Resource Center of the University of Texas Southwestern (UTSW) Medical Center. All protocols were approved by the UTSW Institutional Animal Care and Use Committee. Mice were housed in standard cages that contained three to five mice per cage, with water and standard diet *ad libitum* and a 12 h light/dark cycle. Both male and female mice were analyzed in all experiments. *Nestin-Cre* mice (Stock No. 003771) were obtained from Jackson Laboratory (Bar Harbor, ME) (Tronche et al., 1999). *HoxB7-Cre* and *Ksp-Cre* mice were obtained from O’Brien Kidney Research Core of UT Southwestern. ES cells targeting the second exon of *Tulp3* from EUCOMM (HEPD 0508-5-B01) were used to generate the floxed mice that were obtained from MRC, Harwell as described before (Hwang et al., 2019). For genotyping of *Tulp3^f/f^* mice, we used the following primers: (a) Tulp3-5’Cas-WTF (5’ CCA TTT GTG AGG GTT GCT TT 3’) and Tulp3-Crit-WTR (5’ GCT AAC ACA AGC CCA TGC TA 3’) to detect wild type band (256bp) and floxed band (450 bp), and (b) Tulp3-F1 (5’ AAG GCG CAT AAC GAT ACC AC 3’) and Tulp3-R1 (5’ ACT GAT GGC GAG CTC AGA CC 3’) to detect the deletion. We did not notice any difference between wild type or heterozygous animals for *Tulp3* floxed alleles with or without *Cre* recombinase, and thus were all included as littermate controls as mentioned in respective figure legends. For genotyping *Cre*, following primers were used to detect transgene band: Cre-F (5’ AAT GCT GTC ACT TGG TCG TGG C 3’), Cre-R (5’ GAA AAT GCT TCT GTC CGT TTG C 3’).

### Antibodies and reagents

Anti-Arl13b rabbit polyclonal was from Tamara Caspary (Emory University School of Medicine) (Caspary et al., 2007), Anti-Smo rabbit polyclonal was from Kathryn Anderson (Memorial Sloan Kettering Cancer Center, New York, NY) (Ocbina and Anderson, 2008). Affinity-purified rabbit polyclonal antibody against TULP3 (Mukhopadhyay et al., 2010) and Gpr161 has been described previously (Pal et al., 2016). Anti-Ift139 (Thm1) antibody was from Dr. David Beier (Tran et al., 2008). Commercial antibodies used were against Arl13b (IF, N295B/66 NeuroMab; WB, Proteintech 17711-1-AP), GFP (IF, Abcam ab13970; WB, Abcam ab290), hFAB Rhodamine Anti-Tubulin (Bio-Rad; 12004166), acetylated α-tubulin (mAb 6-11B-1 T6793; Sigma-Aldrich;), S tag (EMD Millipore; MAC112), IRDye 680RD Streptavidin (LI-COR; 926-68079) Myc (Abcam; ab9132), HA (Rat polyclonal;), Flag (Abcam; ab1257;), MBP (New England Biolabs, Inc.), GST (Sigma; G1160) and AC3 (LifeSpan BioSciences; LS-C204505), Aqp2 (1:500 A7310 Sigma rabbit polyclonal; SC515770, Santa Cruz Biotechnology mouse IgG1), Inpp5e (Proteintech; 17797-1-AP), Nphp3 (Proteintech; 22026-1-AP), Lkb1 (Cell Signaling Technologies; 13031T). Fluorescent secondary antibodies for immunofluorescence were from the Jackson Immuno Research Laboratories, Inc., and IRDye 680RD, IRDye 800CW secondary antibodies, and neutravidin 680RD for immunoblotting were from LI-COR Biosciences. Reagents used in this study included PIP Strips Membranes (Thermo Fisher; P23751) and Biotin (Sigma; B4639).

### Plasmids

pG-LAP1 (pCDNA5/FRT/TO-EGFP-TEV-Stag-X) and pG-LAP5 (pEFα-X-Stag-PreScission-EGFP) were from Addgene (Torres et al., 2009). pENTR223-ARL13B fusion construct was from DNASU (HsCD00511796), pENTR221-INPP5E construct was from Life technologies (IOH40212), NPHP3 (encoding 1-203 aa) were synthesized commercially from Geneart (Life Technologies) and Gateway PLUS shuttle clone for CYS1 was from GeneCopoeia (GC-Y0203-CF). N or C terminal LAP tagged retroviral constructs of full length and truncations of TULP3, INPP5E, NPHP3, CYS1 and ARL13B were generated by Gateway cloning into a gatewaytized LAP1 or LAP5 version of pBABE, respectively. Lentiviral constructs of Myc-TULP3 and HA-INPP5E were generated by gateway cloning into pQXIN-Myc and pQXIN-HA destination vectors, respectively. Single or multiple amino acid mutants of TULP3 and ARL13B were generated by Q5 site directed mutagenesis (New England Biolabs). For biotinylation experiments, DEST-pcDNA5-BirA-FLAG N- or C-term destination vectors were from C. Gingras, The Lunenfeld-Tanenbaum Research Institute at Mount Sinai Hospital, Toronto, Canada. Myc tagged full-length and truncated ARL13B fragments were generated by *in vitro* translation using TNT Sp6 high-yield wheat germ protein expression system (Promega; L3261) from pCS2-Myc-ARL13B vectors generated by gateway cloning. FL and N-terminus (1-298 aa) of ARL13B was cloned into pOCC29 (MBP-TEV-GFP-insert-TEV-His) baculovirus vector for insect cell expression (Lemaitre et al., 2019). GST or MBP tagged proteins for bacterial purification were cloned into pGEX or pMalC2 vectors, respectively. GFP-SNAP-ARL13B was generated by cloning SNAP into a BstB1 site of the S-peptide in an N terminal LAP tagged ARL13B retroviral construct (pBABE). A *β*1 Integrin-HaloTag-ARL13B–C-GFP (IA-GFP) was from Takanari Inoue, Johns Hopkins University (Su et al., 2013). The *β*1 Integrin-HaloTag-GFP without ARL13B–C terminus (I-GFP) was generated by Q5 site directed mutagenesis (New England Biolabs).

### Cell culture and generation of stable cell lines

RPE hTERT cells were grown in DMEM F12 media with 10% FBS (Sigma-Aldrich), 0.05 mg/ml penicillin, 0.05 mg/ml streptomycin, and 4.5 mM glutamine. T-Rex-293 (Invitrogen), IMCD3 Flp-In, and Phoenix A (PhA) cells (Indiana University National Gene Vector Biorepository) were cultured in DMEM high glucose (Sigma-Aldrich; supplemented with 10% cosmic serum, 0.05 mg/ml penicillin, 0.05 mg/ml streptomycin, and 4.5 mM glutamine). Transfection of plasmids was done with Polyfect (QIAGEN) or Polyethylenimine (PEI) max. Stable cell lines were generated by retroviral infection or transfection. In many cases, stable lines were flow sorted and further selected for GFP. Control, *Tulp3* ko and *Tulp3^K407I^* MEFs were from E13.5 embryos (Legue and Liem, 2019). Immortalized wildtype, *Arl13b^V358A^* and *Arl13b^hnn^* MEFs were gifts from Tamara Caspary (Gigante et al., 2020).

### Generation of Tulp3 knock out cell lines

CRISPR/Cas9 knockout lines for *Tulp3* were generated in IMCD3 Flp-In (Invitrogen), NIH 3T3 Flp-In and 3T3-L1 (gift of Peter Michaely, UT Southwestern) cells by using guide RNA targeting sequences caccgACGTCGCTGCGAGGCATCTG and caccgTGGCTTTAACCTTCGCAGCC targeting exon 3 in mouse *Tulp3*. Single clones were isolated using serial dilution method. Clonal lines were tested for knockout by Sanger sequencing and immunoblotting for Tulp3.

### Reverse transcription and quantitative PCR

RNA was extracted from paraffin embedded brain sections using deparaffinization silution (Qiagen Cat #19093) and RNA extraction using GenElute mammalian total RNA purification kit (RTN350; Sigma). qRT-PCR was performed with Kicqstart One-Step Probe RT-qPCR ReadyMix (KCQS07; Sigma). Inventoried TaqMan probes for qRT-PCR from Applied Biosystems were used for human *TULP3* and *GAPDH*. Inventoried probes for *Gpr161*, and *Gapdh* were. Reactions were run in CFX96 Real time System (Bio Rad).

### Immunofluorescence of cultured cells and microscopy

Cells were cultured on coverslips until confluent and starved for indicated periods. Cells were fixed with 4% PFA. After blocking with 5% normal donkey serum, the cells were incubated with primary antibody solutions for 1 h at room temperature followed by treatment with secondary antibodies for 30 min along with Hoechst 33342 (Invitrogen). The coverslips were mounted using Fluoromount G (SouthernBiotech). Images were acquired on a microscope (AxioImager.Z1; ZEISS), a sCMOS camera (PCO Edge; BioVision Technologies), and Plan Apochromat objectives (10×/0.45 NA; 20×/0.8 NA; 40×/1.3 NA oil; and 63×/1.4 NA oil) controlled using Micro-Manager software (University of California, San Francisco) at room temperature. Between 8 and 20 z sections at 0.5–0.8-µm intervals were acquired. For quantitative analysis of ciliary localization, stacks of images were acquired from three to eight consecutive fields with confluent cells by looking into the DAPI channel, and percentages of protein-positive ciliated cells were counted. Maximal projections from images of stacks were exported from ImageJ/Fiji (National Institutes of Health) using a custom-written macro (from M. Mettlen, University of Texas Southwestern Medical Center, Dallas, TX) using similar parameters (image intensity and contrast) for image files from the same experiment. For measuring ciliary pixel intensities, image stacks were acquired with z sections at 0.5-µm intervals. An image interval with maximal intensity was chosen, and cilia were demarcated with a region of interest using fluorescence signal for acetylated α-tubulin. The mean pixel intensities for the corresponding protein were exported from ImageJ/Fiji. SNAP experiments were performed as described earlier (Follit and Pazour, 2013; Keppler et al., 2003). Briefly, cells on coverslips were serum starved for 36 hours to induce ciliation. 0.05 µM BG-Block (NEB, S9106) was added for 30 min after which the coverslips were washed in serum starvation media and fixed at different time points. Cells were immunostained with indicated antibodies. 0.3 µM fluorescent SNAP substrate (TMR Star, NEB, S9105) was added along with the primary antibodies

### Tissue processing, immunostaining, and microscopy

Mice were perfused with PBS, and the brains and kidneys were dissected and fixed in 4% paraformaldehyde overnight at 4°C and processed for paraffin embedding and sectioning. For paraffin sectioning, tissues were processed over a 12-hour period using a Thermo-Fisher Excelsior Automated Tissue Processor (A82300001; ThermoFisher Scientific), which dehydrated the tissues through 6 ethanol concentrations, from 50% ethanol to 100% ethanol, cleared through 3 changes of xylene, and infiltrated with wax through 3 Paraplast Plus paraffin baths (39602004; Leica). Samples were embedded in Paraplast Plus using paraffin-filled stainless steel base molds and a Thermo-Shandon Histocenter 2 Embedding Workstation (6400012D; ThermoFisher Scientific). The tissues were then cut in 5 μm thick sections, deparafinned and treated with microwave in Antigen Retrieval citra solution (HK086-9K; BioGenex. Fremont, CA) for 10 min. Sections were then blocked using blocking buffer (1% normal donkey serum [Jackson immunoResearch, West Grove, PA], in PBS) for 1 hour at room temperature. Sections were incubated with primary antibodies against the following antigens; overnight at room temperature or 4C: Acetylated tubulin (1:500, T6793; Sigma mouse IgG2b) Arl13b (1:500, N295B/66; NeuroMab Facility), Aqp2 (1:500 A7310 Sigma rabbit polyclonal; SC515770, Santa Cruz Biotechnology mouse IgG1), Inpp5e (1:500 17797-1-AP; Proteintech), Nphp3 (1:500 Proteintech; 22026-1-AP) or Lkb1 (1:500 Cell Signaling Technologies; 13031T). After three PBS washes, the sections were incubated in secondary antibodies (Alexa Fluor 488-, 555-, 594-, 647-conjugated secondary antibodies, or anti-mouse IgG isotype-specific secondary antibodies; 1:500; Life Technologies, Carlsbad, CA or Jackson ImmunoResearch). Cell nuclei were stained with DAPI (Sigma) or Hoechst 33342 (Life technologies). Slides were mounted with Fluoromount-G (0100-01; Southern Biotech) and images were acquired with a Zeiss AxioImager.Z1 microscope or a Zeiss LSM780 confocal microscope.

### Proximity biotinylation experiments

T-Rex-293 cells were co-transfected with 5–7.5 µg each of LapN-TULP3 or LapN-TULP3 mutants/fragments and pDEST-pcDNA5-BirA-FLAG N/C-term expressing ARL13B or the CD8 linker-BirA or FibrocystinCLS-BirA fusion controls. The media was supplemented with 20 µM biotin for 8-12 hours after transfection. Cells were harvested using PBS with 2 mM EDTA and 2 mM EGTA 48 h after transfection. Cells were lysed by resuspending and nutating for 20 min in 50 mM Tris-HCl, pH 7.4, 200 mM KCl, 1 mM MgCl_2_, 1 mM EGTA, 10% glycerol, 1 mM DTT, 0.6% IGEPAL CA-630, 1 mM AEBSF, and 0.01 mg/ml each of leupeptin, pepstatin, and chymostatin. Lysates were centrifuged at 12,000 *g*for 10 min followed by tandem IPs (Fig. 5). In brief, the GFP immunoprecipitates were first digested with TEV protease for 16 h at 4°C. The supernatants were subjected to secondary IP with S protein agarose. The resulting secondary IPs were analyzed by Western blotting. Blots were probed with antibodies against S tag (mouse monoclonal MAC112) and Flag (goat polyclonal ab1257) followed by visualization using IRDye-tagged secondary antibodies. IRdye-tagged streptavidin was used for confirming biotinylation signal on the blots.

### *In vitro* binding experiments

GST tagged proteins (20 μg) were incubated with 30 μL packed volume of glutathione sepharose beads in LAP150N (50 mM HEPES pH 7.4, 150 mM KCl, 1 mM EGTA, 1 mM MgCl_2_, 10% glycerol, 0.05% NP-40) buffer at 4°C for 2-4 hours. Beads were washed with LAP150N three times and then incubated with increasing amounts of IVT Myc-ARL13B (2.5, 5 and 10 uL of the IVT reaction) or insect cell purified GFP-ARL13B (2.5, 5 and 10 ug) in LAP100N (50 mM HEPES pH 7.4, 150 mM KCl, 1 mM EGTA, 1 mM MgCl_2_, 10% glycerol, 0.05% NP-40) or LAP150N buffer at 4°C or room temperature respectively for 1 hour. In figure 5C, 50 ug of the indicated MBP proteins were added along with IVT Myc-ARL13B. Flowthroughs were collected, beads were washed, eluted by boiling in SDS sample buffer and subjected to western blotting.

### PIP and membrane lipid blots

Binding of recombinant proteins to pre-spotted PIP and membrane lipid strips (Thermo Fisher; P23751) was performed according to (Dowler et al., 2002). After blocking strips in Licor blocking buffer for 1 hr at room temperature, the strips were incubated in Licor blocking buffer containing 1 μg/ml of recombinant GST, GST-TULP3 or GST-TULP3^K389I^ protein overnight at 4°C. After washing three times in TBS-T, the blots were immunoblotted for GST.

### Mass spectrometry

In the Taplin proteomics core (^LAP^TULP3^K389I^), excised gel bands were cut into approximately 1 mm^3^ pieces. For proximity biotinylation experiments, CD8-Gpr161 IC3^WT^-BirA*, CD8-Gpr161 IC3^5A^-BirA*, CD8-MCHR1 IC3^WT^-BirA*, CD8-MCHR1 IC3^5A^-BirA* were co-transfected with ^LAP^TULP3 in TREx-293 cells. After LAP tandem affinity purification, excised gel bands were submitted for post translational analysis at Taplin proteomics core for biotinylated peptides. Gel pieces were then subjected to a modified in-gel trypsin digestion procedure (Shevchenko et al., 1996). Gel pieces were washed and dehydrated with acetonitrile for 10 min followed by removal of acetonitrile. Pieces were then completely dried in a speed-vac. Rehydration of the gel pieces was with 50 mM ammonium bicarbonate solution containing 12.5 ng/μl modified sequencing-grade trypsin (Promega, Madison, WI) at 4°C. After 45 min, the excess trypsin solution was removed and replaced with 50 mM ammonium bicarbonate solution to just cover the gel pieces. Samples were then placed in a 37°C room overnight. Peptides were later extracted by removing the ammonium bicarbonate solution, followed by one wash with a solution containing 50% acetonitrile and 1% formic acid. The extracts were then dried in a speed-vac (∼1 h). The samples were then stored at 4°C until analysis. On the day of analysis, the samples were reconstituted in 5-10 μl of HPLC solvent A (2.5% acetonitrile, 0.1% formic acid). A nano-scale reverse-phase HPLC capillary column was created by packing 2.6 µm C18 spherical silica beads into a fused silica capillary (100 µm inner diameter x ∼30 cm length) with a flame-drawn tip (Peng and Gygi, 2001). After equilibrating the column each sample was loaded via a Famos auto sampler (LC Packings, San Francisco CA) onto the column. A gradient was formed, and peptides were eluted with increasing concentrations of solvent B (97.5% acetonitrile, 0.1% formic acid). As peptides eluted, they were subjected to electrospray ionization and then entered an LTQ Orbitrap Velos Pro ion-trap mass spectrometer (Thermo Fisher Scientific, Waltham, MA). Peptides were detected, isolated, and fragmented to produce a tandem mass spectrum of specific fragment ions for each peptide. Peptide sequences (and hence protein identity) were determined by matching protein databases with the acquired fragmentation pattern by the software program, Sequest (Thermo Fisher Scientific, Waltham, MA) (Eng et al., 1994). Post translational analysis was performed using probability-based scoring (Beausoleil et al., 2006). All databases include a reversed version of all the sequences and the data was filtered to between a one and two percent peptide false discovery rate.

### Statistical Analyses

Statistical analyses were performed using Student’s *t*-test for comparing two groups or Tukey’s post hoc multiple comparison tests between all possible pairs using GraphPad Prism. Nonparametric Matt Whiney-U test were performed for intensity plots using GraphPad Prism.

## Acknowledgements

This project was funded by PKD Foundation postdoctoral fellowship to VRP (214F19a), the National Institutes of Health (1R01GM113023 to S.M. and NS097928 to K.L.), a PKD Foundation grant to K. L. (232G18) and a Cancer Prevention and Research Institute of Texas grant (RR170063 to J.B.W), We thank molecular pathology and mass spectrometry cores and mouse animal care facility in UT Southwestern. We thank Ross Tomaino (Taplin mass spectrometry facility) for assistance with mass spectrometry. We acknowledge kind gifts of reagents from Tamara Caspary, Kathryn Anderson, David Beier, Peter Jackson, Peter Michaely, Takanari Inoue and Anne-Claude Gingras. We thank Issei Shimada and John Shelton for *Nestin-Cre* brain sections. We thank Sandii Constable for comments on the manuscript.

## Author contributions

V. P. and S. M conceived the project, designed experiments, analyzed most of the data, and wrote the paper with inputs from all authors. S. H. generated cell lines and performed immunofluorescence experiments, B. N. S. performed kidney immunofluorescence experiments, H. B. generated mass spectrometry data on biotinylated samples, V. M.T. and J. B. W. provided help with insect cell purifications and reagents, E.L and K. L. generated *Tulp3* mutant MEFs.

---

**Figure S1.**
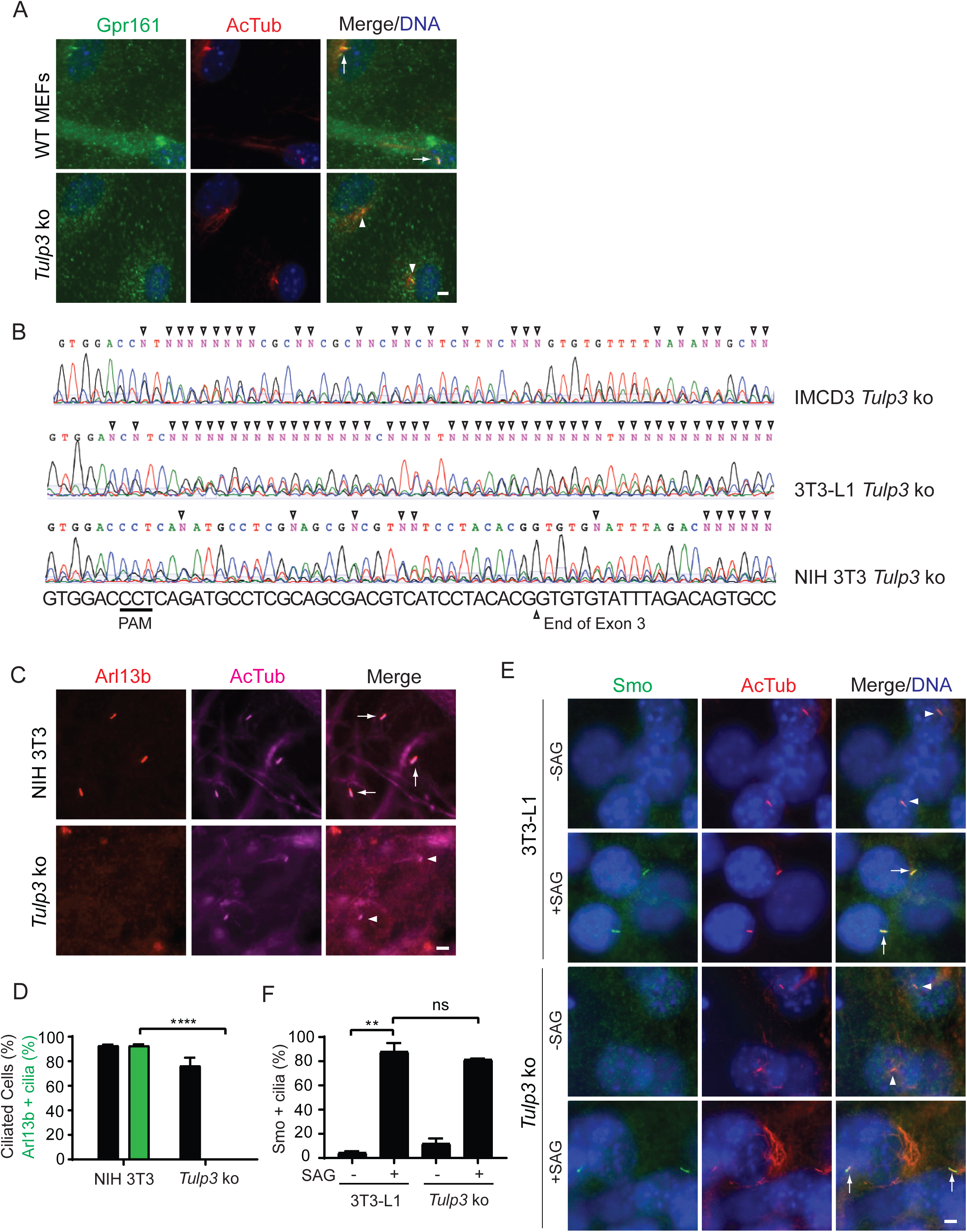
(Related to Figure 1). Tulp3 determines ciliary trafficking of Arl13b. **(A)** MEFs from wild type or *Tulp3* knockout (ko) mice were serum starved upon confluence for 24 h before fixation. Fixed cells were immunostained for Gpr161 (green) along with acetylated tubulin (AcTub, red) and counterstained for DNA (blue). Quantification shown in Fig. 1B. **(B)** Genomic DNA was isolated from *Tulp3* ko 3T3-L1, NIH 3T3 or IMCD3 and sequenced using exon3 specific primers. Open arrowheads point to mutations. N, non-specific nucleotide. **(C-D)** Wildtype and *Tulp3* ko NIH 3T3 cells were grown to confluency and starved further for 48 h to promote ciliation before fixing. The fixed cells were immunostained for Arl13b (green) along with acetylated tubulin (red) and counterstained for DNA. Total counted cells are >200 for each condition. Data represent mean ± SD. **(E-F)** Wild type or *Tulp3* knock out 3T3-L1 cells were grown to confluency and further cultured for 72 hours to promote ciliation. The cells were treated with 500 nM Smo agonist (SAG) for 24 hours to induce Smo ciliary localization before fixing. Immunostaining was performed for Smo, Acetylated tubulin and DNA. Arrows indicate cilia positive for Smo while arrow heads indicate Smo negative cilia. Smo positive cilia from B were counted from two experiments (E), and >100 cilia were counted for each condition. Data represent mean ± SD. Scale, 5 μm. ****, p<0.0001; ns, not significant. Arrows indicate cilia positive for the indicated proteins (Gpr161 (A) or Arl13b (C) or Smo (E)), while arrow heads indicate negative cilia.

**Figure S2.**
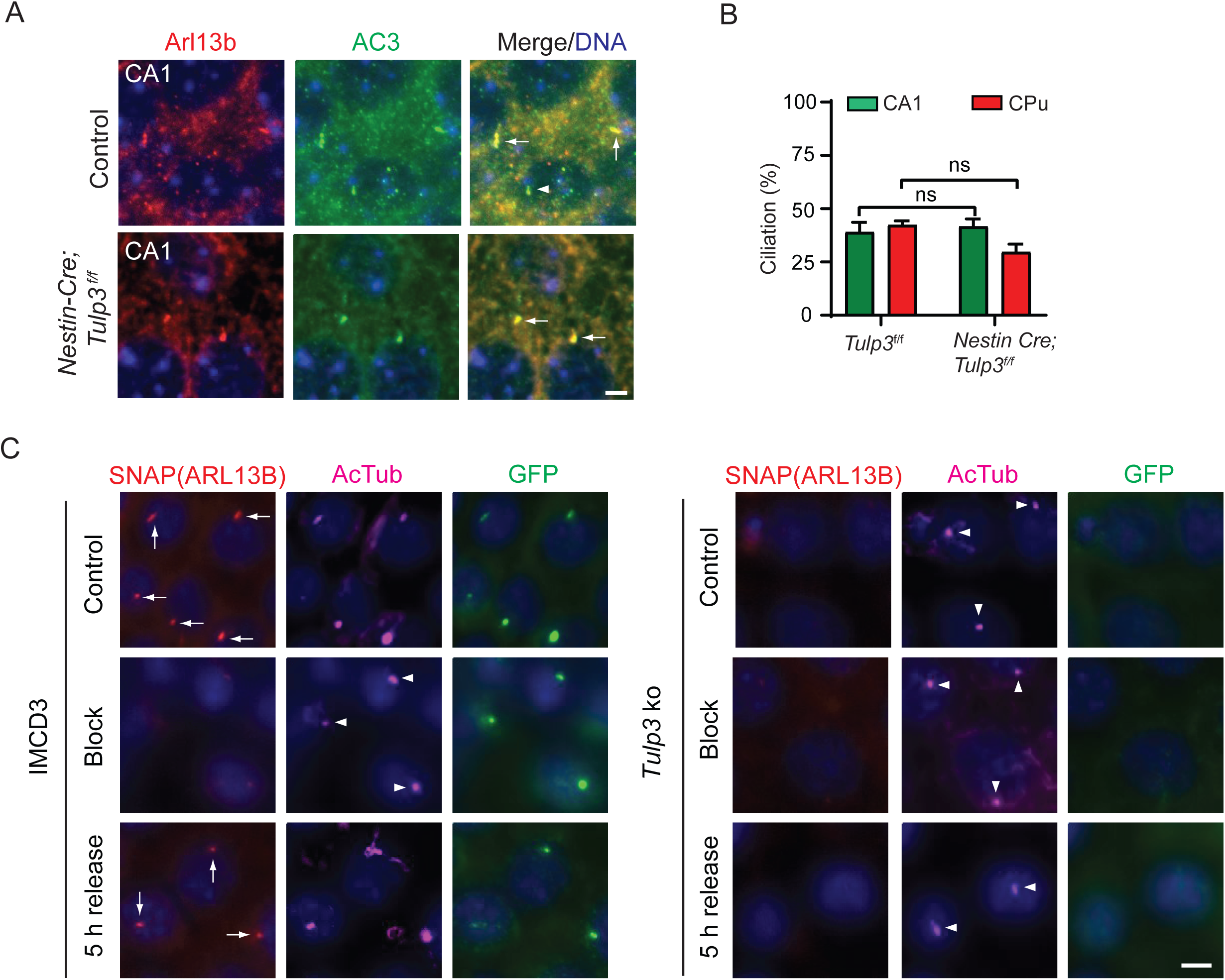
(related to Figure 1). Ciliary localization of Arl13b was unaffected in *Nestin-Cre*; *Tulp3^f/f^* brain regions. **(A)** Brain sections from CA1 regions in hippocampus from either control or *Nestin-Cre*; *Tulp3*^f/f^ mice at P7 were immunostained for Arl13b and AC3 and counterstained for DNA. Quantification shown in Figure 1H. **(B)** Cilia (AC3) positive cells were counted from CA1 regions in hippocampus and caudate-putamen (CPu) regions of brains from either control or *Nestin-Cre; Tulp3^f/f^* mice. Total counted cells were >200 from one mouse for control and two mice for *Nestin-Cre; Tulp3^f/f^*. Data represent mean ± SD. **(C)** Representative images for Figure 1J. Scale, 5 μm. ns, not significant.

**Figure S3.**
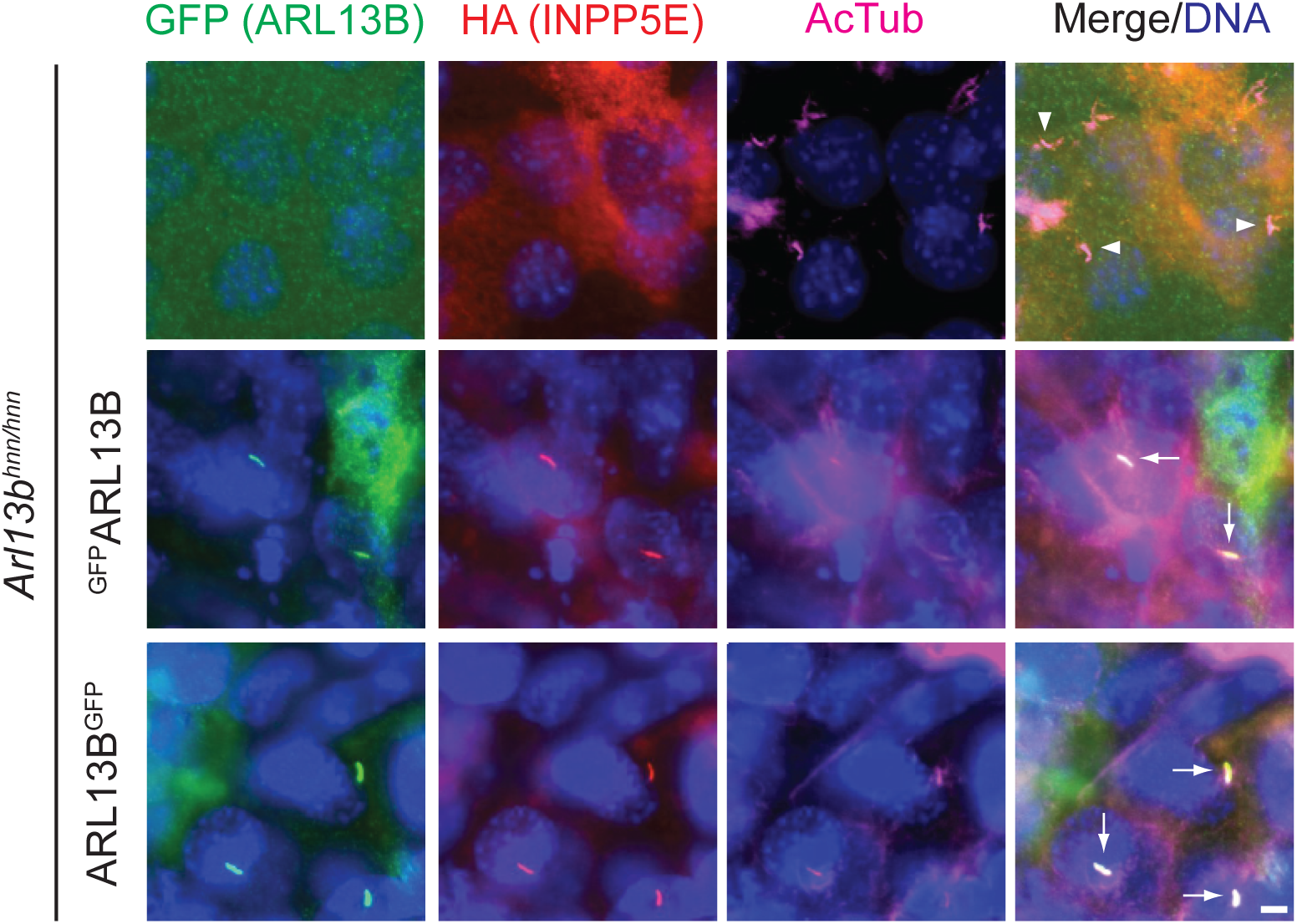
(Related to Figure 2). Tulp3 determines ciliary trafficking of Arl13b dependent cargoes. N term- or C term-GFP-Stag (LAP) tagged ARL13B stably expressed in immortalized *Arl13b^hnn^* MEFs along with ^HA^INPP5E were starved for 48 h and fixed. Fixed cells were immunostained for GFP (ARL13B), HA (INPP5E), acetylated tubulin (AcTub) and counterstained for DNA. Both N- or C-tagged ARL13B fusions trafficked to cilia in *Arl13b^hnn^* background. ^HA^INPP5E levels in cilia were also restored by either fusion, suggesting rescue from their stable expression. Scale, 5 μm.

**Figure S4.**
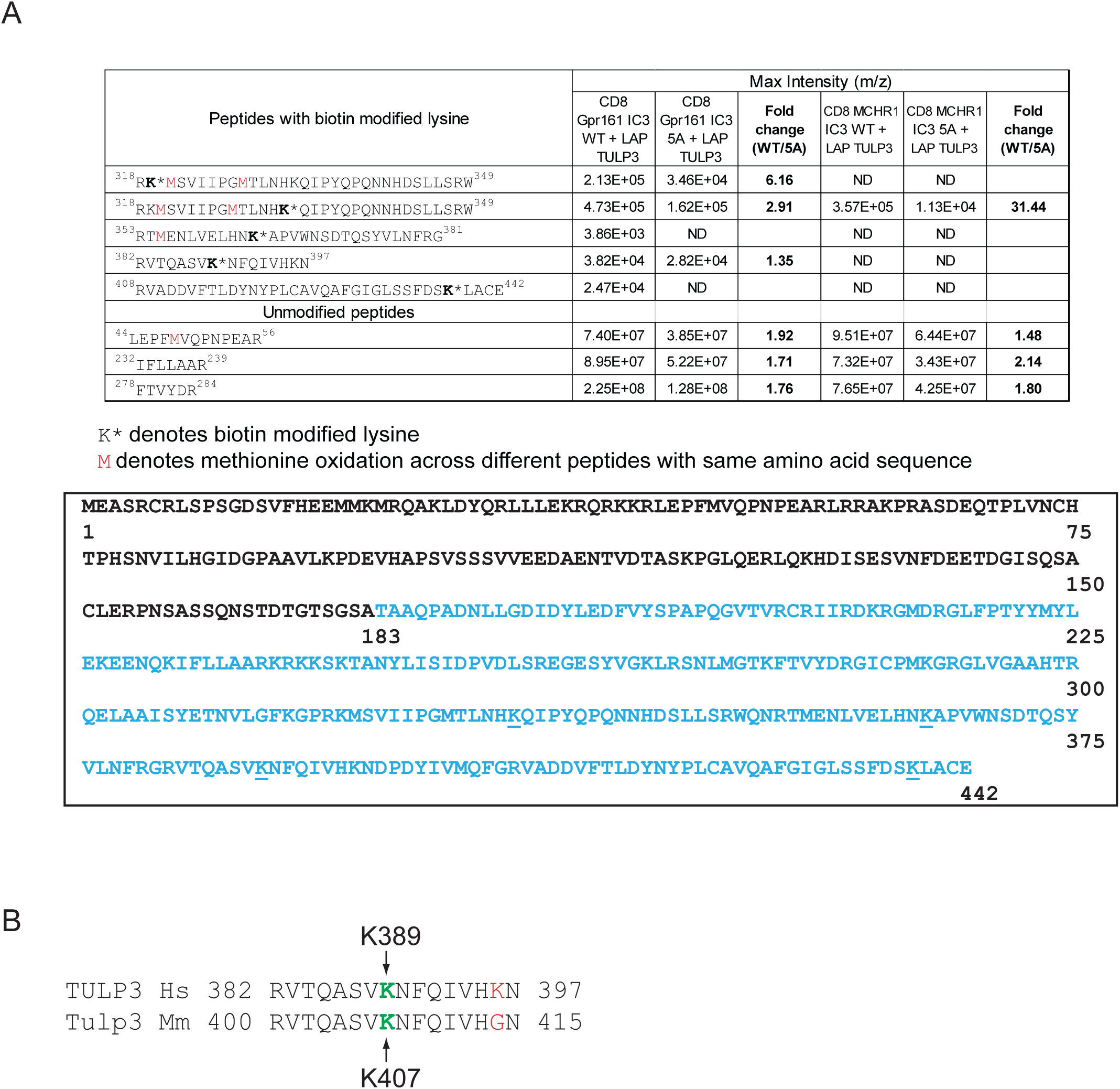
(Related to Figure 6): Lysine K389 in TULP3 is required for interaction with its cargoes. **(A)** Sites in TULP3 biotinylated by ciliary localization sequences of GPCRs Gpr161 and MCHR1 in comparison to corresponding mutated CLSs that are non-ciliary (**Figure S4A**) (Badgandi et al., 2017) identified by mass spectrometry (see Methods). **(B)** Alignment of tubby region of TULP3 flanking the Lysine K389 in human TULP3 (Hs, *Homo sapiens* NP_003315.2) with that from mouse (Ms, *Mus musculus* NP_035787.1).

**Figure S5.**
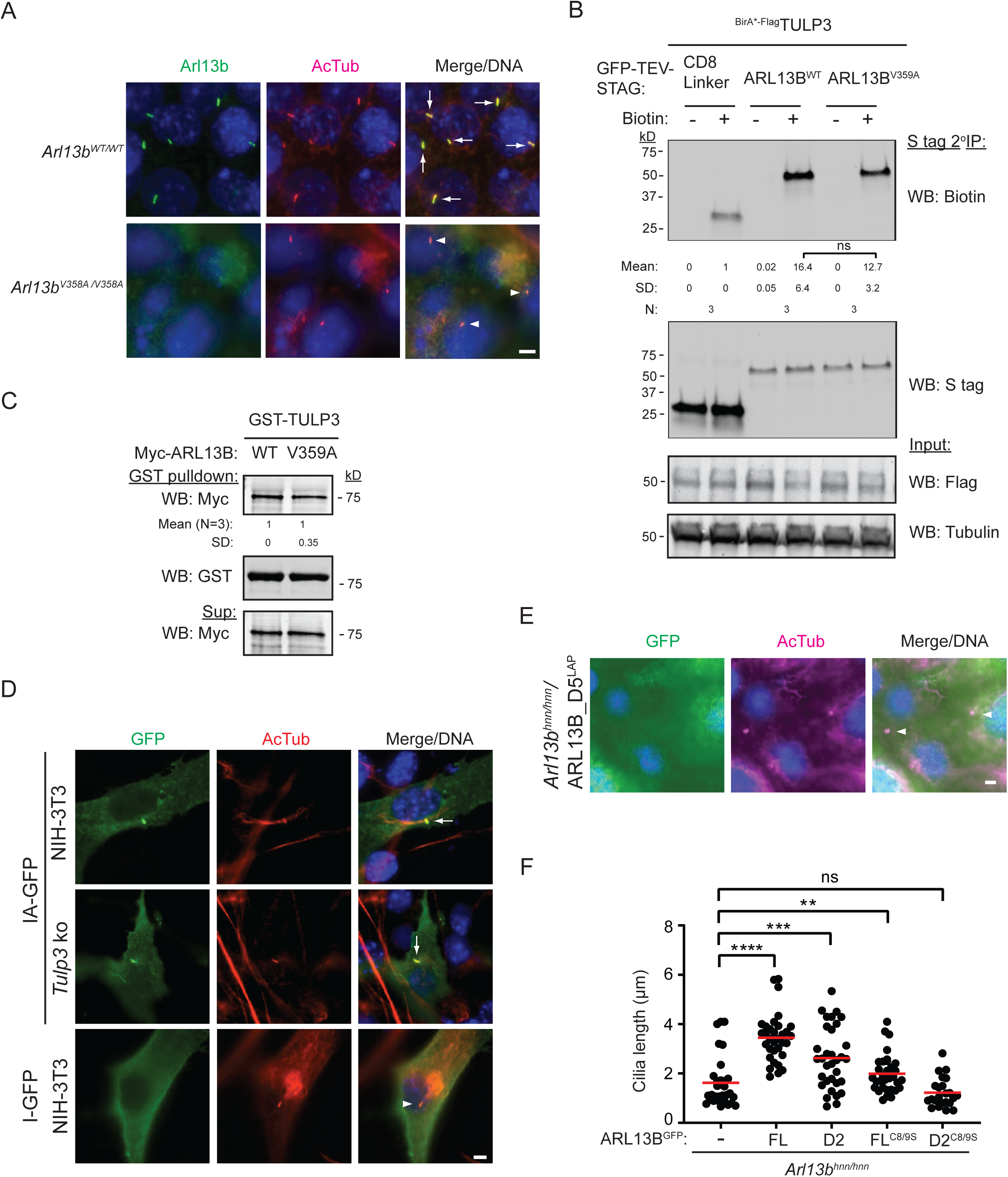
(Related to Figure 8): Arl13b domains required for ciliary localization. **(A)** Immortalized *Arl13b^V358A/V35A^* MEFs were starved for 48 h and immunostained for Arl13b, acetylated tubulin (AcTub) and counterstained for DNA. **(B)** T-Rex 293 cells were co-transfected with BirA* tagged TULP3 along with LAP tagged ARL13B or V359A mutant and processed as shown in Figure 5A. Mean ± SD values indicate Biotin/S-tag ratios normalized to CD8 inker control. “n” indicates the number of experiments performed **(C)** Bacterially purified ^GST^TULP3 protein bound to glutathione Sepharose beads was tested for binding with *in vitro* translated (IVT) Myc full length ARL13B and *V359A* mutant as in Figure 8B. **(D)** NIH 3T3 wildtype or *Tulp3* ko cells were transfected with IA-GFP or I-GFP (with deleted ARL13B C-terminus from IA-GFP) and starved for 48 h. Fixed cells were immunostained for GFP and AcTub. **(E)** *Arl13b^hnn^* cells stably expressing C-LAP tagged D5 were serum starved for 48 h before fixing and immunostained for GFP, acetylated tubulin (AcTub) and counterstained for DNA. **(F)** Cilia lengths of cell lines as described in 8(E). Only ciliated cells in the *Arl13b^hnn^* background stably expressing HA-tagged INPP5E were counted. *Arl13b^hnn^* cells are about ∼20% ciliated (Larkins et al., 2011). Scale, 5 μm. ****, p<0.0001; ***, p<0.001; **, p<0.01; ns, not significant. Arrows indicate cilia positive for the indicated proteins while arrowheads indicate negative cilia.

## Supplementary Text

Different reports on ciliary targeting sequences in ARL13B report contrasting results. Here we describe some of these results in chronological order.

First, by transfecting N-terminal (up to GTPase domain) or C-terminal fragment (from CC domain) in wild type mammalian cells, it was shown that neither construct localized to cilia (Hori et al., 2008). The N-terminal region (up to GTPase domain) also showed self-association. Please note that the N-terminal fragment that we found to be ciliary included the CC domain (**Figure 8A**, D2). Arl13b mutants lacking CC domain (18-202) in *Chlamydomonas* completely reduced Arl3-GEF activity (Gotthardt et al., 2015) and could prevent access to the amphipathic helix.

Second, by injecting *Arl13b* mRNA into 1-4 cell stage zebrafish embryos, multiple regions were shown to be necessary for ciliary localization (Duldulao et al., 2009). These included part of the GTPase domain, the coiled coil motif following the GTPase domain and the C-terminus of the protein starting from the coiled coil domain. Many of the ciliary localization and rescue experiments were carried out in a knockout background. Most importantly, a D2 like construct (1-307 aa that included N-terminus beyond the CC domain) was found to be rescuing body curvature and cyst formation despite not being ciliary in localization. We find the D2 construct to be rescuing ^HA^INPP5E levels in cilia but also cilia localized, but only partially rescuing cilia lengths in *Arl13b^hnn^* MEFs compared to wild type (**Figure 8A, E**).

Third, initial reports in *C. elegans* suggested a role of ARL13B palmitoylation in redistributing the protein to the cell body of phasmid neurons, but proximal ciliary localization persisted (Cevik et al., 2010; Cevik et al., 2013). Deleting the C-terminus RVxP motif (RVVP sequence in *C. elegans*) also did not prevent proximal ciliary localization of ARL13B in phasmid neurons, rather caused it to distribute more distally in addition to forming cell body aggregates (Cevik et al., 2013). These ARL13B fusions were all expressed in presence of wild type protein. Of note, the RVVP motif deleted ARL13B transgene functions as a dominant negative when injected in wild type *C. elegans* for dye-filling (such dye filling defects are also a feature of *ARL13B* null allele) (Cevik et al., 2013). Such dominant negative effects suggest mutual interactions in compromising wild-type protein function. Similar dominant negative effects are also seen in palmitoylation mutants (Cevik et al., 2010), suggesting coupling with membrane to be critical for mutual interactions in compromising wild-type protein function.

Fourth, by transiently transfecting N-terminal (1-212, disrupting CC) and C-terminal Arl13b (210-427) in *Arl13b^hnn^* background, it was shown that neither fragment was ciliary (Larkins et al., 2011). Please note that the N-terminal construct that we found to be ciliary included the CC domain (**Figure 8A**, D2). A single amino acid mutation in mouse RVxP motif, *Arl13b^V358A^*, was shown to be nonciliary upon transient transfection in *Arl13b^hnn^* MEFs (Mariani et al., 2016). Palmitoylation motif mutants (*Arl13b^C8/9S^*) were also found to be partially ciliary (∼20% compared to ∼80% localization for wild type) when transfected in *Arl13b^hnn^* background (unlike our results using stable expression that showed robust localization in cilia and ^HA^INPP5E rescue in cilia) (**Figure 8A, 8E**). MEFs generated from endogenous *Arl13b^V358A/358A^* knock-in embryos also showed no steady state levels of the mutant protein in cilia (Gigante et al., 2020) (**Figure S5A**).

Fifth, another study using transfected tagged *ARL13B* constructs in wild type ARL13B background reported the RVEP-to-AAEA mutation to disrupt ciliary localization in RPE cells, but also form aggregates in the cytoplasm (Nozaki et al., 2017). GTPase domain mutants (T35N, R79Q) were ciliary as reported earlier (Humbert et al., 2012). The INPP5E binding region in ARL13B was pinpointed to both GTPase and CC domains (Nozaki et al., 2017) and GTPase domain mutants (T35N, R79Q) also reduced INPP5E binding (Humbert et al., 2012; Nozaki et al., 2017). Deletion of only GTPase domain (unlike zebrafish experiments (Duldulao et al., 2009)) or swapping an ARF6 GTPase domain still retained ciliary localization (Nozaki et al., 2017). The authors were not able to infer if ARL13B(ΔCC) localizes to cilia, as this mutant fused to tags formed aggregates in the cytoplasm (Nozaki et al., 2017), and could explain lack of ciliary expression seen in zebrafish experiments (Duldulao et al., 2009).

## KEY RESOURCE TABLE

| REAGENT OR RESOURCE | SOURCE | IDENTIFIER |
| --- | --- | --- |
| **Antibodies and related** |  |  |
| Mouse monoclonal anti-Acetylated tubulin | Sigma | Cat# T6793<br>RRID: AB_477585 |
| Rabbit polyclonal anti-Arl13b (IF) | Tamara Caspary (Emory University) |  |
| Rabbit polyclonal anti-Arl13b (WB) | Proteintech | Cat# 17711-1-AP<br>RRID: AB_2060867 |
| Mouse monoclonal anti-Arl13b (IF) | NeuroMab Facility | Cat# N295B/66<br>RRID: AB_2750771 |
| Chicken anti-GFP (IF) | Abcam | Cat# ab13970<br>RRID: AB_300798 |
| Rabbit polyclonal anti-Smo | Kathryn Anderson (MSKCC) | (Ocbina and Anderson, 2008) |
| Rabbit polyclonal anti-Tulp3 | Peter Jackson (Stanford University) | (Mukhopadhyay et al. 2010) |
| Rabbit polyclonal anti-Gpr161 | Custom-made | (Pal et al. 2016) |
| Rabbit polyclonal anti-IFT139 | David Beier (Seattle Children's) | (Tran et al., 2008) |
| Rabbit polyclonal anti-GFP (WB) | Abcam | Cat# ab290<br>RRID: AB_303395 |
| Mouse monoclonal anti-S tag | EMD Millipore | Cat# MAC112 |
| Goat polyclonal anti-Myc tag | Abcam | Cat# ab9132<br>RRID: AB_307033 |
| Rat monoclonal anti-HA | Roche | Cat# 11867423001<br>RRID: AB_390918 |
| Goat polyclonal anti-DDDDK (Flag) tag | Abcam | Cat# ab1257<br>RRID: AB_299216 |
| Mouse monoclonal anti-MBP | NEB | Cat# E8032S |
| Mouse monoclonal anti-GST | Sigma | Cat# G1160<br>RRID: AB_259845 |
| Rabbit polyclonal anti-ADCY3 | LSBio | Cat# LS-C204505 |
| Rabbit polyclonal anti-Aqp2 | Sigma | Cat# A7310<br>RRID: AB_476762 |
| Mouse monoclonal anti-Aqp2 | Santa Cruz | Cat# sc-515770<br>RRID: AB_2810957 |
| Rabbit polyclonal anti-INPP5E | Proteintech | Cat# 17797-1-AP<br>RRID: AB_2167120 |
| Rabbit polyclonal anti-NPHP3 | Proteintech | Cat# 22026-1-AP<br>RRID: AB_2878975 |
| Rabbit Monoclonal anti-LKB1 | CST | Cat# 13031T |
| hFAB Rhodamine anti-Tubulin | Bio-Rad | Cat# 12004165<br>RRID: AB_2884950 |
| IRDye 680 Streptavidin | LI-COR | Cat# 926-68079 |
| Alexa Fluor 488-, 594-, 647- conjugated secondary antibodies | Life Technologies |  |
| IRDye tagged secondary antibodies | LI-COR |  |
| Hoechst 33342 | Life Technologies | H3570 |
| BG-Block | NEB | S9106 |
| TMR Star | NEB | S9105 |
| **Chemicals, Media, and Kits** |  |  |
| Penicillin/Streptomycin | Sigma | P4333 |
| Glutamine | Sigma | G7513 |
| Polyfect | Qiagen | 301105 |
| PEI-Max | Polysciences | 24765-2 |
| DMEM-High Glucose | Sigma | D5796 |
| DMEM F12 | Sigma | D6421 |
| Bovine calf serum | Sigma | 12133C |
| Fetal bovine serum | Sigma | F0926 |
| Cosmic serum |  |  |
| Paraformaldehyde | Electron microscopy solutions | 15710 |
| Normal donkey serum | Jackson Immuno Research, West Grove, PA | Cat# 017-000-121<br>RRID: AB_2337258 |
| Fluoromount-G | Southern Biotech | 0100-01 |
| Permunt | ThermoFisher Scientific | SP15-100 |
| Biotin | Sigma | Cat# B4639 |
| GenElute mammalian total RNA purification kit | Sigma | RTN350 |
| DNase I | Sigma | D5307 |
| SYBR Green Quantitative RT-qPCR Kit | Sigma | QR0100 |
| Kicqstart One-Step Probe RT-qPCR ReadyMix | Sigma | KCQS07 |
| TNT Sp6 high-yield wheat germ protein expression system | Promega | L3261 |
| Synthetic mounting media | Fisher Chemical | SP15 |
| Mini-PROTEAN TGX Precast Protein Gels | Bio-Rad |  |
| **Experimental Models: Organisms** |  |  |
| Mouse: *Tulp3<sup>fl/fl</sup>* | MRC, Harwell | (Hwang et al, 2019) |
| Mouse: *HoxB7-Cre* | O'Brien Kidney Research Core of UT Southwestern |  |
| Mouse: *Ksp-Cre* | O'Brien Kidney Research Core of UT Southwestern |  |
| Mouse: *Nestin-Cre* | JAX | Stock No. 003771 |
| **Experimental Models: Cell Lines** |  |  |
| IMCD3 Flp-In | Peter Jackson (Stanford University) | (Torres et al, 2009) |
| RPE hTERT | ATCC | ATCC CRL-4000 |
| NIH 3T3 Flp-In | ATCC | R76107 |
| 3T3-L1 | Peter Michaely<br>(UT Southwestern) |  |
| TRex-293 FLP-In | Life Technologies | R78007 |
| **Experimental Models: Primary cells** |  |  |
| *Tulp3* knockout MEFs | This study |  |
| *Tulp3* K407I/K407I MEFs | This study |  |
| **Experimental Models: Immortalized MEFs** |  |  |
| *Arl13b*<sup>hnn</sup> MEFs and wild type controls | Tamara Caspary<br>(Emory University) | (Larkins et al, 2011) |
| *Arl13b*<sup>V358A</sup> MEFs and wild type controls | Tamara Caspary<br>(Emory University) | (Gigante et al, 2020) |
| **Recombinant DNA** |  |  |
| pGLAP5 | Addgene | 19706 |
| pG-LAP1 | Addgene | 19702 |
| pENTR223-ARL13B | DNAsu | HsCD00511796 |
| pENTR221-INPP5E | Life Technologies | IOH40212 |
| NPHP3 (1-203 aa) | GeneArt | Synthesized |
| Gateway PLUS shuttle clone for CYS1 | GeneCopoeia | GC-Y0203-CF |
| Gatewaytized LAP1 or LAP5 in pBABE | This study |  |
| pQXIN-Myc-TULP3 | This study |  |
| pQXIN-HA-INPP5E | This study |  |
| pcDNA5-BirA-FLAG N-DEST | Anne-Claude Gingras |  |
| pcDNA5-BirA-FLAG C-DEST | Anne-Claude Gingras |  |
| pCS2-Myc-DEST |  |  |
| pOCC29 |  | (Lemaitre et al., 2019) |
| GFP-SNAP-ARL13B in pBABE | This study |  |
| **Software and Algorithms** |  |  |
| ImageJ software | National Institutes of Health, Bethesda, MD |  |
| GraphPad Prism | GraphPad, La Jolla, CA |  |
| PictureFrame 3.0 acquisition software | Optronics, Inc. Goleta, CA, USA |  |
| Sequest | Thermo Fisher Scientific, Waltham, MA |  |
| **Other** |  |  |
| PIP Strip | Thermo Fisher | P23751 |
| Thermo-Fisher Excelsior Automated Tissue Processor | ThermoFisher Scientific | A82300001 |
| Paraplast Plus paraffin bath | Leica | 39602004 |
| Thermo-Shandon Histocenter 2 Embedding Workstation | ThermoFisher Scientific | 6400012D |
| Leica stereomicroscope (M165 C) with digital camera (DFC500) | Leica |  |
| Zeiss stereomicroscope (Discovery.V12) | Zeiss |  |
| Zeiss LSM780 confocal microscope | Zeiss |  |
| Leica DM2000 photomicroscope | Leica |  |
| Sakura DRS-601 x-y-z robotic-stainer | Sakura-FineTek,<br>Torrance, CA | DRS-601 |
| CFX96 thermocycler | Bio-Rad |  |
| Superfrost® Plus slides | Fisher Scientific | 12-550-15 |
| Vitros 250 chemistry analyzer | GMI Inc., Ramsey, MN |  |

